# Alveolar differentiation drives resistance to KRAS inhibition in lung adenocarcinoma

**DOI:** 10.1101/2023.09.29.560194

**Authors:** Zhuxuan Li, Xueqian Zhuang, Chun-Hao Pan, Yan Yan, Rohit Thummalapalli, Jill Hallin, Stefan Torborg, Anupriya Singhal, Jason C. Chang, Eusebio Manchado, Lukas E. Dow, Rona Yaeger, James G. Christensen, Scott W. Lowe, Charles M. Rudin, Simon Joost, Tuomas Tammela

## Abstract

Lung adenocarcinoma (LUAD), commonly driven by *KRAS* mutations, is responsible for 7% of all cancer mortality. The first allele-specific KRAS inhibitors were recently approved in LUAD, but clinical benefit is limited by intrinsic and acquired resistance. LUAD predominantly arises from alveolar type 2 (AT2) cells, which function as facultative alveolar stem cells by self-renewing and replacing alveolar type 1 (AT1) cells. Using genetically engineered mouse models, patient-derived xenografts, and patient samples we found inhibition of KRAS promotes transition to a quiescent AT1-like cancer cell state in LUAD tumors. Similarly, suppressing *Kras* induced AT1 differentiation of wild-type AT2 cells upon lung injury. The AT1-like LUAD cells exhibited high growth and differentiation potential upon treatment cessation, whereas ablation of the AT1-like cells robustly improved treatment response to KRAS inhibitors. Our results uncover an unexpected role for KRAS in promoting intra-tumoral heterogeneity and suggest targeting alveolar differentiation may augment KRAS-targeted therapies in LUAD.

**Significance:** Treatment resistance limits response to KRAS inhibitors in LUAD patients. We find LUAD residual disease following KRAS targeting is composed of AT1-like cancer cells with the capacity to reignite tumorigenesis. Targeting the AT1-like cells augments responses to KRAS inhibition, elucidating a therapeutic strategy to overcome resistance to KRAS-targeted therapy.

## Introduction

*KRAS* is the most commonly mutated proto-oncogene across all human cancers. KRAS is a prominent oncogenic driver in three of the most lethal cancers – non-small cell lung cancer (NSCLC), colorectal cancer, and pancreas cancer (1,2). Thus, KRAS has long represented one of the most attractive therapeutic targets in oncology (1). *KRAS* mutations are particularly prevalent in the lung adenocarcinoma (LUAD) subtype of NSCLC, which is driven by mutant KRAS in ∼30% of cases (3). 41% of *KRAS* mutations in LUAD involve a conversion of glycine in position 12 to cysteine [*KRAS(G12C)*] (3). Recent exciting developments have led to the FDA approval of the first direct inhibitors of KRAS(G12C) in LUAD patients (3). Despite the promise of KRAS(G12C) inhibitors, early results from human clinical studies indicate that only 30-50% of patients respond (4). Furthermore, the duration of response in these patients is limited by acquired resistance involving activation of parallel signaling pathways that bypass KRAS, lineage transformation, or drug desensitizing mutations in *KRAS* itself (5,6). A next generation of mutant KRAS allele-specific inhibitors is emerging, including KRAS(G12D) inhibitors (1,7). However, early results suggest that patient-derived xenograft (PDX) models established from LUAD patients may be less responsive to these inhibitors than pancreas cancer models (7). These preclinical and clinical data suggest that LUAD may harbor lineage-specific features that limit the efficacy of KRAS inhibitors.

LUAD predominantly arises from alveolar type 2 (AT2) epithelial cells, which function as facultative alveolar stem cells by self-renewing and replacing alveolar type 1 (AT1) cells. Following injury, AT2 cells proliferate and expand in response to receptor tyrosine kinase (RTK) ligands emanating from the lung stroma (8). Upon cessation of these mitogenic signals, a subset of AT2 cells differentiates into AT1 cells, which completes alveolar re-epithelialization (8). Oncoproteins mimic these mitogenic signals but, unlike physiological mitogens, they are perpetually active in LUAD cells. This relentless oncogene activity conspires with acquired genetic and epigenetic changes to drive the emergence of tumors composed of diverse phenotypically distinct cancer cell states (9). Among many aspects of malignant progression, the clinical relevance of this intra-tumoral heterogeneity manifests in the distinct capacities of cancer cell states for treatment resistance. Specific cell states driving drug resistance have been described in e.g. colon (10), basal skin tumors (11), and glioblastomas (12). In most cases, the resistant states are characterized by features of normal tissue stem cells or progenitors (10-13). A recent report described a progenitor-like cell state that was enriched in LUAD residual disease following EGFR-or ALK-targeted therapy (14). The effect of targeting oncogenic KRAS on cancer cell states in LUAD has not been investigated.

Given that efficacy of KRAS inhibitors in patients is limited by intrinsic and acquired resistance poses several important questions: What is the cellular source of cancer cells that acquire resistance to KRAS inhibitors? Are there LUAD cell states that are intrinsically independent of KRAS signaling and, if such cell states exist, can targeting them eradicate resistance to KRAS inhibition? To address these questions, we employed genetically engineered mouse models (GEMMs) of LUAD, based on the activation of oncogenic ***<u>K</u>****ras(G12D)* and deletion of **<u>p</u>**53 (*“**KP**”* model) in AT2 cells. These models recapitulate key molecular and histopathological features of LUAD tumors in humans (15), including responses to chemotherapy (16).

## Results

### Targeting Kras promotes an AT1-like differentiation program in LUAD cells

To investigate the impact of targeting KRAS in autochthonous LUAD tumors, we generated two germline *KP* models that enable doxycycline (Dox)-inducible tumor-specific short hairpin RNA (shRNA)-mediated silencing of *Kras*; we also generated a control shRNA model targeting *Renilla* luciferase (**Fig. 1A**) (17,18). In these mice, an shRNA is coupled to a GFP cDNA, placed downstream of a tetracycline-responsive (*TET*) promoter, and targeted via recombinase-mediated cassette exchange to the ubiquitously expressed *Col1a1* locus (**Supplementary Fig. 1A-C**) (19,20). In this *KP; RIK; TET-GFPshRNA* system, adenoviral delivery of Cre recombinase under the control of an AT2 cell-specific surfactant protein C promoter [AdSPC-Cre (21)] leads to the expression of oncogenic *Kras^G12D^*and deletion of the tumor suppressor p53, resulting in LUAD development. In addition, the delivery of Cre recombinase activates the expression of reverse tetracycline transactivator-3 (rtTA3) and the mKate2 fluorescent protein (***<u>r</u>****tTA3-**<u>I</u>**RES-m**<u>K</u>**ate2*, or ***RIK*** cassette) in the *KP* mutant cells. Upon systemic administration of Dox to the mice, rtTA3 will bind to tetracycline responsive elements (TRE) and the expression of a GFP-linked shRNA targeting *Kras* (*shKras247* or *shKras462*) or *Renilla* (control) specifically in the LUAD cells ensues (**Fig. 1B**).

**Figure 1.**
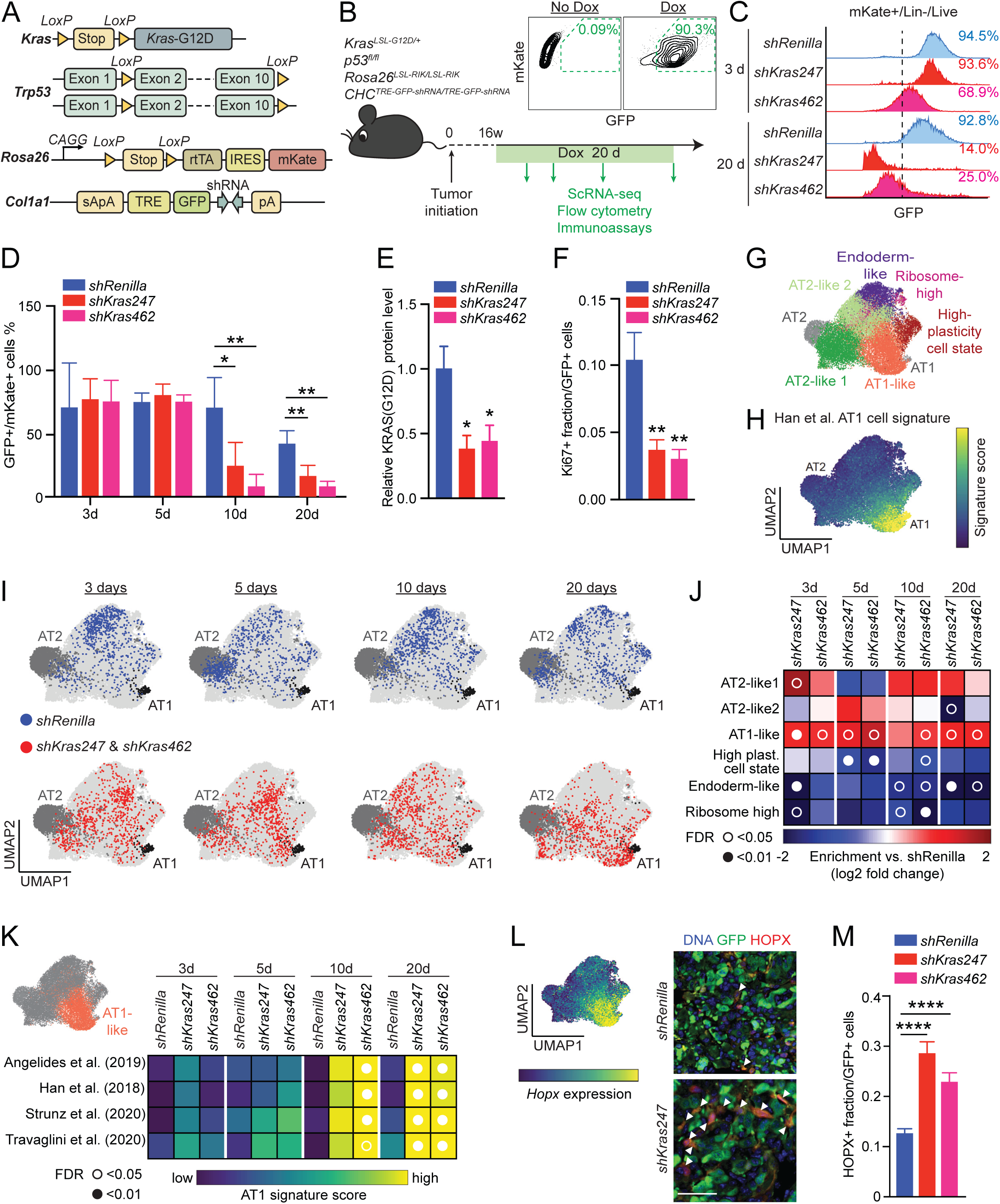
Targeting Kras promotes an AT1-like differentiation program in LUAD cells. (**A**) Genetically engineered *KP; RIK; TET-GFPshRNA* mouse model enabling doxycycline (Dox)- inducible shRNA expression in *KP* LUAD cells. (**B**) Experimental design for targeting *Kras* using the *KP; RIK; TET-GFPshRNA* system. Inset: Flow cytometry plot showing induction of GFPshRNA in the *KP* tumors *in vivo.* Cells gated as CD45^-^/CD31^-^/CD11b^-^/F480^-^/TER119^-^/DAPI^-^ (live). (**C**) GFP expression in mKate^+^/ CD45^-^/CD31^-^/CD11b^-^/F480^-^/TER119^-^/DAPI^-^ (live) cells in *shRenilla* and *shKras* groups at different time points. (**D**) Proportion of GFP^+^/mKate^+^ cells in total mKate^+^ LUAD cell pool in *shRenilla* and *shKras* groups at indicated time points; *n* = 3 replicates per model for each time point. (**E**) Relative KRAS(G12D) protein expression in isolated primary GFP^+^/mKate^+^ *KP; RIK; TET-GFPshRNA* LUAD cells after 20 days on Dox; *n* = 3 mice/group. (**F**) Fraction of proliferating (Ki67^+^) GFP^+^/mKate^+^ LUAD cells after 5 days on Dox; *n* ≥ 13 mice/group. (**G**) Unsupervised clustering of GFP^+^/mKate^+^ single LUAD cell transcriptomes, colored and annotated based on Marjanovic et al. (9). Normal healthy AT2 and AT1 single-cell transcriptomes (gray) isolated from wild-type mice are co-embedded. (**H**) Projection of wild-type mouse AT1 cell gene expression signature (47) onto the UMAP space shown in (G). (**I**) Location of *KP; RIK; TET-GFPshRNA* LUAD cell transcriptomes in the UMAP space following expression of *shRenilla* control or *shKras247*/*shKras462* at the indicated time points; *n* = 3-4 mice/group (750 randomly sampled cells per condition). (**J**) Fold change (log2) in the proportion of the distinct cancer cell subsets shown in (G) following expression of *shKras* at the indicated time points. Open circle: p < 0.05; closed circle: *p* < 0.01 (*t* test vs. *shRenilla* with individual tumors as biological replicates). (**K**) Signature score of four independent AT1 cell signatures in the AT1-like LUAD cell state (orange) in each model. Shown is the average score of the single-cell transcriptomes per group. Note the increasing expression of AT1 cell genes over time. Open circle: *p* < 0.05; closed circle: *p* < 0.01 (*t* test vs. *shRenilla* with individual tumors as biological replicates). (**L**) *Left*: projection of *Hopx* expression level onto the UMAP in (G). *Right*: representative images of HOPX immunofluorescence (red) in the LUAD cells expressing the indicated *shRNAs* (green). Arrowheads indicate HOPX^+^/GFP^+^ cells. Scale bar: 100 µm. (**M**) Quantification of the fraction of HOPX^+^/GFP^+^ cells within the total GFP^+^ cell pool; *n* ≥ 36 tumors/experimental group. Two-way ANOVA was used in (D), (E), (F), and (M) to test for statistical significance: **** *p* < 0.0001; ** *p* < 0.01; * *p* < 0.05. Error bars indicate SEM.

To test these models, we induced LUAD in the *KP; RIK; TET-GFPshRNA* mice using AdSPC-Cre. We isolated mKate^+^ cancer cells from all three models at 16 weeks post-tumor initiation (PTI), a time point when the tumors had progressed to fully formed adenocarcinoma (21), and generated adherent cell lines (**Supplementary Fig. 1D**). Induction of the shRNAs targeting *Kras* depleted KRAS protein expression, suppressed ERK phosphorylation, and inhibited growth of these cell lines, whereas induction of the *shRenilla* control had no effect (**Supplementary Fig. 1E, F**). To evaluate the shRNAs *in vivo*, *KP; RIK; TET-GFPshRNA* mice were placed on Dox for 3, 5, 10, and 20 days starting at 16 weeks PTI (**Fig. 1B**). The proportion of GFP^+^/mKate2^+^ LUAD cells was consistent in all shRNA models at 3 and 5 days following Dox exposure. However, after 10 or 20 days of shRNA expression, the fraction of GFP^+^/mKate2^+^ cells was significantly lower in the *shKras247* and *shKras462* mice than in the *shRenilla* controls (**Fig. 1C, D**), indicating selection against *Kras* knockdown. This selection likely occurs via silencing of the *TET-GFPshRNA* allele, which has been widely reported (17). We observed efficient depletion of the mutant KRAS(G12D) protein in GFP^+^/mKate^+^ LUAD cells expressing *shKras247* and *shKras462* when compared to *shRenilla* control *in vivo* (**Fig. 1E; Supplementary Fig. 1G**). Induction of either *Kras* shRNA suppressed the proliferation of the *KP* cancer cells *in vivo* and *in vitro* when compared to *shRenilla* control (**Fig. 1F; Supplementary Fig. 1H**). Finally, *Kras* knockdown promoted apoptosis of the LUAD cells *in vivo*, which was most prominent at 3 days of Dox exposure (**Supplementary Fig. 1I**). Taken together, these results establish our TET-inducible shRNA systems as a powerful genetic model of KRAS inhibition.

To investigate the impact of targeting KRAS on LUAD cell state heterogeneity, we examined the GFP^+^/mKate2^+^ cells that retained shRNA expression at 3, 5, 10, or 20 days of Dox exposure by single-cell mRNA sequencing (scRNA-seq) (**Fig. 1B**). The single-cell expression profiles spanned six clusters with distinct expression patterns by unsupervised clustering (**Fig. 1G-I; Supplementary Data Table 1**). These clusters corresponded to cell states that we had previously identified in fully formed *KP* LUAD tumors (**Fig. 1G**) (9). Interestingly, the trajectories of the LUAD cells expressing *shKras247* or *shKras462* began to separate from the *shRenilla* control at 3 days, culminating in a dramatic phenotypic shift at 20 days following shRNA expression (**Fig. 1I**). Surprisingly, this shift was defined by the enrichment of the LUAD cells predominantly in a defined region in phenotypic space exhibiting AT1-like identity (**Fig. 1H-J**). Conversely, the expression of *shRenilla* control over the 20-day treatment period had no effect on the topology of the phenotypic space occupied by cancer cells (**Fig. 1I**).

The LUAD cells in the AT1-like cluster distinctly expressed genes associated with AT1 differentiation (**Fig. 1H**). This signature showed some similarity to a previously published targeted therapy resistance signature observed in EGFR mutant LUAD cells (**Supplementary Fig. 2A**) (22). Notably, the *shRenilla* control tumors also harbored a subset of AT1-like LUAD cells, but the fraction of cells in this state is smaller (**Fig. 1I, J**) and the correlation of these cells with published AT1 gene expression signatures was less significant than in the *shKras* tumors (**Fig. 1K**). This indicates that – even in the AT1-like state – targeting *Kras* pushes LUAD cells further along the AT1 differentiation trajectory. Among the most prominent AT1 marker genes induced by *Kras* silencing was *Hopx* (**Fig. 1L**). We observed robust expression of HOPX protein in cancer cells in *shKras* tumors, but not in *shRenilla* tumors (**Fig. 1L-M; Supplementary Fig. 2B, C**). Notably, despite the expression of AT1 markers, the LUAD cells did not exhibit morphological features of AT1 cells, such as thinning and spreading on basement membranes (8), in response to KRAS inhibition.

### AT1-like LUAD cells are quiescent and exhibit low MAPK activity

We hypothesized that the AT1-like cancer cells persisting after KRAS suppression may be less dependent on KRAS activity. To investigate this, we evaluated phosphorylation of the KRAS effector ERK as a surrogate marker for KRAS signaling activity in unperturbed *KP* LUAD tumors and found highly variable pERK signaling activity in the cancer cells (**Fig. 2A**). Interestingly, HOPX^+^ AT1-like LUAD cells showed very low levels of pERK when compared to HOPX^-^ LUAD cells (**Fig. 2B, C**). Furthermore, we found the AT1-like LUAD cells exhibited low *Kras* expression and significantly diminished KRAS downstream transcriptomic output compared to the other LUAD cell states (**Supplementary Fig. 2D, E**). In line with lower KRAS signaling activity, we observed the HOPX^+^ LUAD cells were rarely positive for the proliferation marker Ki67 (**Fig. 2D, E**), suggesting that they are quiescent. These results suggest that the AT1- like differentiation state may be less dependent on KRAS than the other LUAD cell states, which may explain resistance of this state to KRAS inhibition.

**Figure 2.**
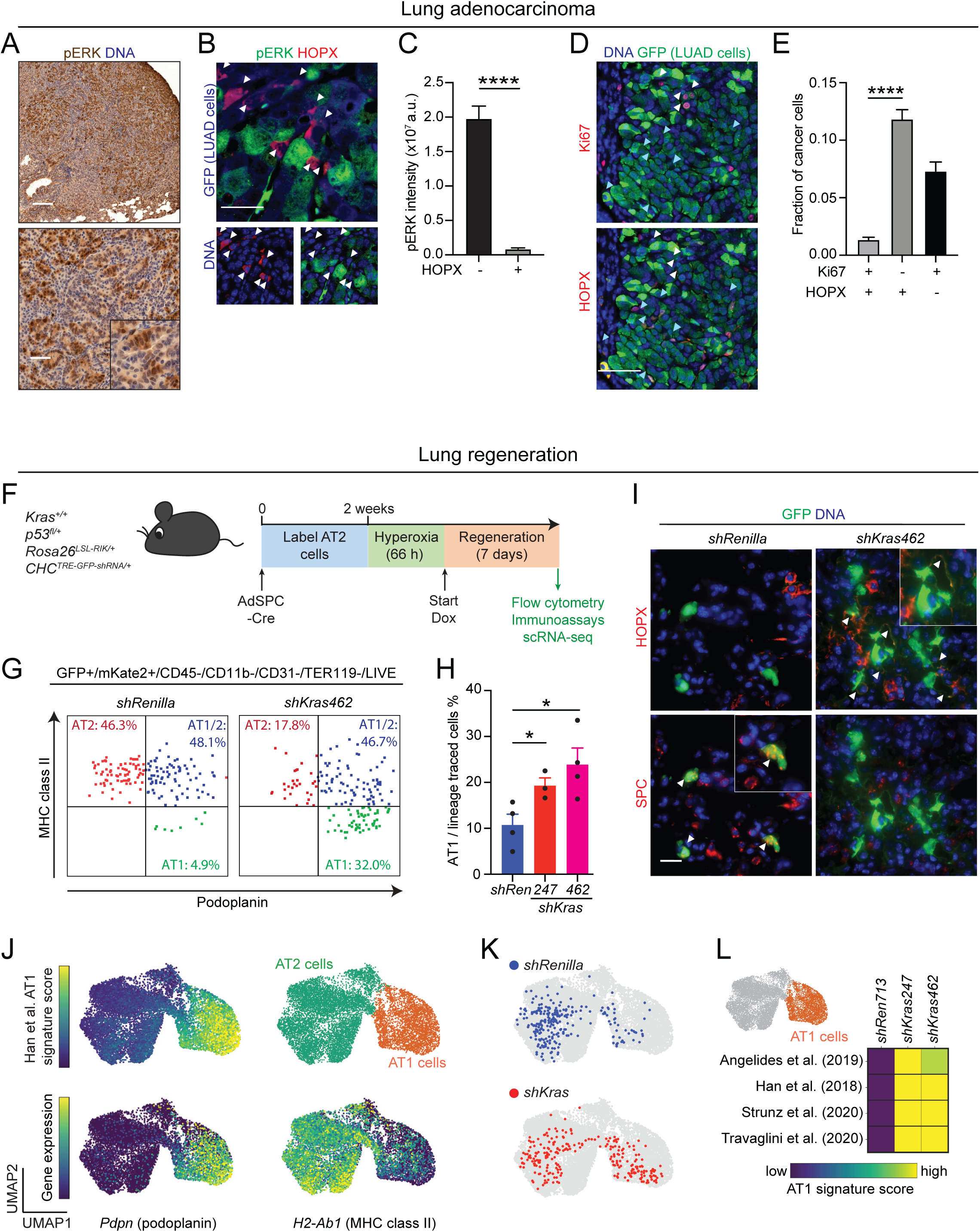
KRAS suppresses AT1 differentiation in LUAD and in the regenerating lung. (**A**) Immunohistochemical staining for pERK (brown) in an autochthonous *KP* LUAD tumor at 16 weeks post-initiation. Scale bars: 200 µm (top) and 100 µm (bottom). (**B**) Representative images showing pERK (green) and HOPX (red) immunofluorescence in a *KP; RIK; TET-GFPshRenilla* LUAD tumor at 16 weeks post-tumor initiation + 5 days on Dox. GFP (blue in top image) marks LUAD cells expressing the *shRenilla* control *shRNA*. Note absence of pERK immunoreactivity in the HOPX^+^/GFP^+^ cells (arrowheads) Scale bar: 50 µm. (**C**) Quantification of pERK staining intensity in HOPX^+^/GFP^+^ vs. HOPX^-^/GFP^+^ LUAD cells in *KP; RIK; TET-GFPshRenilla* tumors at 16 weeks post-tumor initiation + 5 days on Dox; *n* = 168 tumors. An unpaired *t* test was used to test significance. (**D**) Representative images showing Ki67^+^ (red in top image) and HOPX^+^ (red in bottom image) in GFP^+^ LUAD cells in *KP; RIK; TET-GFPshRenilla* LUAD tumors at 16 weeks post-tumor initiation + 5 days on Dox. Note mutual exclusivity of HOPX (turquoise arrowheads) and Ki67 (white arrowheads) staining. Scale bar: 50 µm. (**E**) Quantification of the fraction of Ki67^+^/HOPX^+^/GFP^+^, Ki67^-^HOPX^-^/GFP^+^, and Ki67^+^/HOPX^+^/GFP^+^ LUAD cells in *KP; RIK; TET-GFPshRenilla* LUAD tumors at 16 weeks post-tumor initiation + 5 days on Dox; *n* = 59 tumors. (**F**) Outline of experiment combining AT2 cell lineage-tracing and perturbation of wild-type *Kras* using the *RIK; TET-GFPshRNA* system. (**G**) Representative flow cytometry plots depicting expression of the AT2 marker MHC class II (*y*-axis) and AT1 marker podoplanin (*x*- axis) in *RIK; TET-GFPshRenilla* vs. *RIK; TET-GFPshKras462* mice following lineage-tracing, hyperoxia injury, and 7-day exposure to Dox. (**H**) Quantification of proportion of AT1 cells within total pool of lineage-traced AT2 cells; *n* = 3-5 mice/group. (**I**) Immunofluorescence for HOPX (red, top row) and SPC (red, bottom row). White arrowheads indicate HOPX^+^ AT1 cells in the top row, SPC^+^ AT2 cells in the bottom row. Scale bar: 20 µm. (**J**) Single-cell transcriptomes of lineage-traced (GFP^+^/mKate2^+^) *RIK; TET-GFPshRenilla* vs. *RIK; TET-GFPshKras462* cells following hyperoxia injury, and 7-day exposure to Dox, co-embedded with primary AT2 and AT1 cell transcriptomes isolated from wild-type uninjured lungs. Heatmaps indicate AT1 gene expression score or *H2-ab1* (MHC class II) or *Pdpn* (podoplanin) gene expression; unsupervised clustering separates AT2 (green) and AT1 (orange) cells. (**K**) Lineage-traced cells isolated from *RIK; TET-GFPshRenilla* (blue) or *RIK; TET-GFPshKras462* (red) mice are projected into UMAP space (250 random samples cells per condition). (**L**) Heatmap showing four previously published healthy AT1 cell signatures in the AT1 space (orange) in the indicated *RIK; TET-GFPshRNA* models. Shown is the average score over all single-cell transcriptomes per group; *n* = 2-4 models/group. Two-way ANOVA was used in (E), and (H) to test for statistical significance: **** *p* < 0.0001; * *p* < 0.05. Error bars indicate SEM.

### KRAS suppresses AT1 differentiation during lung regeneration

In homeostasis and regeneration of the healthy lung, AT1 cells arise from AT2 cells (8). Similarly, we found targeting KRAS promotes AT1 differentiation and quiescence of LUAD cells. This tantalizing analogy prompted us to examine the role of KRAS in AT2/AT1 differentiation during lung regeneration leveraging our *RIK; TET-GFPshRNA* system in *Kras* wild-type mice. We poised the TET-inducible *shRNA* system in AT2 cells of these mice using AdSPC-Cre and induced hyperoxia injury, followed by induction of the *shRNAs* targeting *Kras* or *Renilla* control (**Fig. 2F**). Silencing *Kras* led to a significant increase in the proportion of traced cells expressing the AT1 markers podoplanin and HOPX (**Fig. 2G-I**). Furthermore, targeting *Kras* promoted the acquisition of AT1 transcriptomic features (**Fig. 2J-L; Supplementary Data Table 2**), indicating that suppressing KRAS robustly promotes AT2 differentiation towards the AT1 lineage.

### Enrichment of AT1-like LUAD cells in residual disease following KRAS inhibition

We next sought to investigate the impact of pharmacologic inhibition of oncogenic KRAS on LUAD cell states. To do this, we treated mice bearing autochthonous *KP; Rosa26^tdTomato/+^* (*KPT*) tumors with the KRAS(G12D) allele-specific inhibitor MRTX1133 starting at 16 weeks PTI for 5 or 20 days (**Fig. 3A**). We observed a significant reduction in tumor burden and cancer cell proliferation as well as increased apoptosis in the MRTX1133 treated mice compared to vehicle controls (**Fig. 3B-E**). Similar to the shRNA model, MRTX1133 promoted the enrichment of LUAD cells expressing the AT1 marker HOPX (**Fig. 3F**) and induced AT1 gene expression programs (**Fig. 3G-J**). The extent of AT1 lineage gene expression was most profound at 20 days following KRAS inhibition with MRTX1133 (**Fig. 3I, J**). Notably, in contrast to the *shKras* models, MRTX1133 also enriched for a cell cluster mapping within an AT2-like cancer cell state (**Fig. 3G-I**). Furthermore, transcriptomic changes induced by KRAS inhibition showed similarity to receptor tyrosine kinase (RTK) inhibitor resistance signatures observed previously in LUAD cells (**Supplementary Fig. 3A**) (14,22). Conversely, cisplatin chemotherapy did not induce AT1 differentiation in the autochthonous *KP* LUAD tumors (**Supplementary Fig. 3B-D**), suggesting that alveolar differentiation may be a unique resistance mechanism to therapies targeting the RTK-KRAS axis.

**Figure 3.**
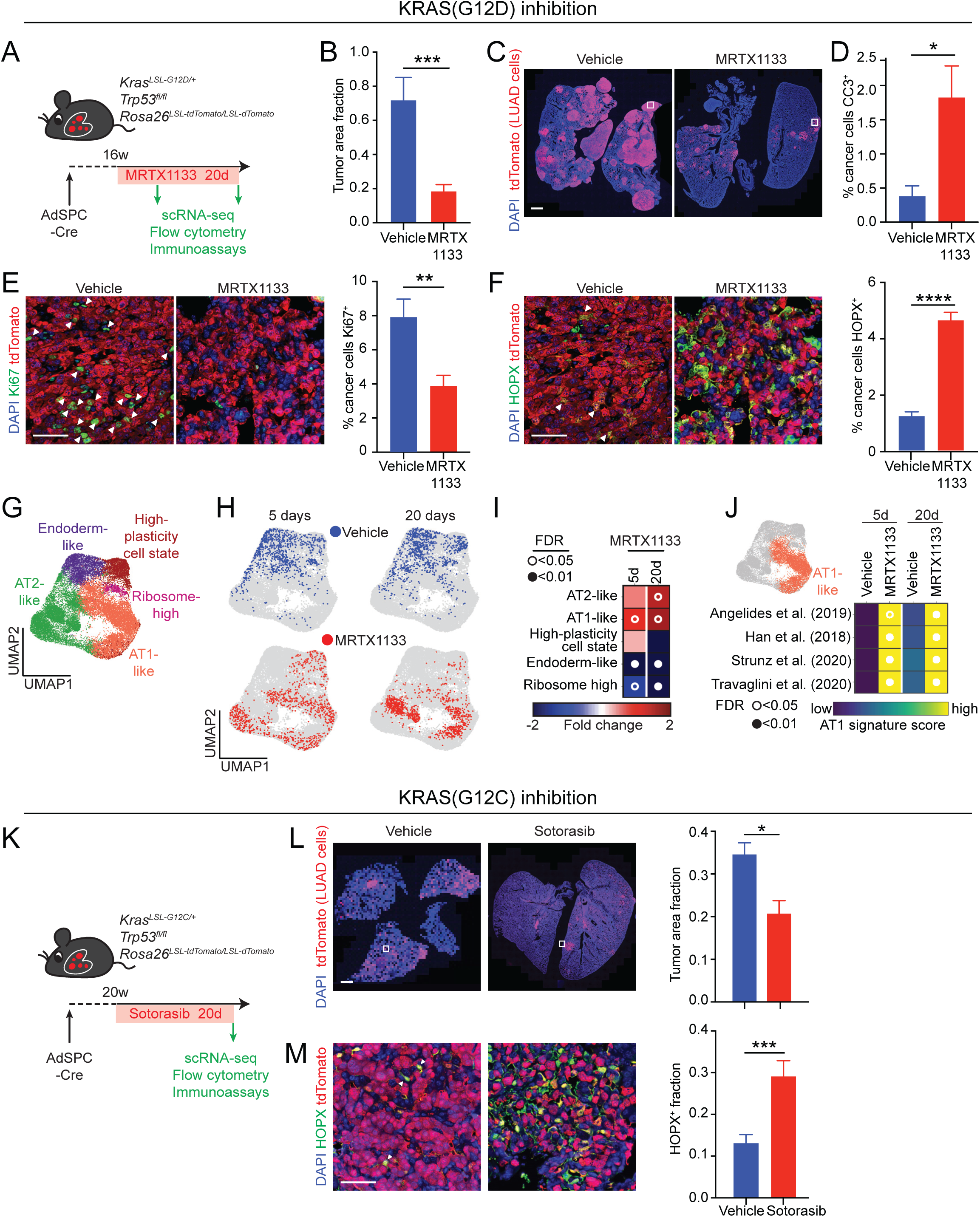
Pharmacologic allele-specific inhibition of KRAS promotes an AT1-like differentiation program in LUAD cells. (**A**) Outline of experimental design to investigate KRAS(G12D) inhibition in autochthonous *KPT* LUAD tumors using MRTX1133. (**B**) Quantification of tumor burden (tdTomato+ area/total lung cross-sectional area) at 16 weeks post-tumor initiation + 20 days on MRTX1133 or vehicle control; *n* ≥ 6. (**C**) Representative images of tdTomato immunofluorescence in *KPT* lung tumors after 20 days of MRTX1133 or vehicle administration. Scale bar: 1 mm. (**D**) Quantification of apoptotic (cleaved caspase 3 [CC3]^+^/tdTomato^+^) cancer cells at 16 weeks post-tumor initiation + 5 days on MRTX1133 or vehicle control (*n* ≥ 17). (**E**) *Left*: Representative images of Ki67 and tdTomato immunofluorescence magnified from the boxed region in (C). Scale bar: 50 µm. *Right*: Quantification of proliferating (Ki67^+^/tdTomato^+^) cancer cells in LUAD tumors at 16 weeks post-tumor initiation + 20 days on MRTX1133 or vehicle control (*n* ≥ 37). (**F**) *Left*: Representative images of HOPX and tdTomato immunofluorescence in the boxed region in (C). Scale bar: 50 µm. *Right*: Quantification of AT1-like (HOPX^+^/tdTomato^+^) cancer cells in LUAD tumors at 16 weeks post-tumor initiation + 20 days on MRTX1133 or vehicle control (*n* ≥ 37). (**G**) Unsupervised clustering of tdTomato^+^/CD45^-^/CD31^-^/CD11b^-^/F480^-^/TER119^-^/DAPI^-^ single LUAD cell transcriptomes, colored and annotated based on Marjanovic et al. (9). (**H**) Location of the LUAD cell transcriptomes in the UMAP space following exposure to vehicle control (blue) or MRTX1133 (red) at the indicated time points (*n* = 3-4 mice/group). (**I**) Fold change in the proportion of the distinct cancer cell subsets shown in (C) following MRTX1133 therapy or vehicle control at the indicated time points. Open circle: *p* < 0.05; closed circle: *p* < 0.01 (*t* test vs. vehicle with individual tumors as biological replicates). (**J**) Signature score of four independent healthy AT1 cell signatures in the AT1-like LUAD cell state (orange) following MRTX1133 therapy or vehicle control. Shown is the average score over all single-cell transcriptomes per group. Note increasing expression of AT1 cell genes over time. Open circle: *p* < 0.05; closed circle: *p* < 0.01 (*t* test vs. vehicle with individual tumors as biological replicates). (**K**) Outline of experimental design to investigate KRAS(G12C) inhibition in autochthonous *KRAS(G12C)PT* LUAD tumors. (**L**) *Left*: Representative images of tdTomato immunofluorescence in *KRAS(G12C)PT* lung tumors at 16 weeks post-tumor initiation + 20 days on sotorasib or vehicle. Scale bar: 1 mm. *Right*: Quantification of surface tumor burden following 20 days of MRTX1133 therapy or vehicle control; *n* ≥ 6. (**M**) *Left*: Representative images of HOPX and tdTomato immunofluorescence in the boxed region in (L). Scale bar: 50 µm. Right: Quantification of AT1-like (HOPX^+^/tdTomato^+^) cancer cells in LUAD tumors at 16 weeks post-tumor initiation + 20 days on sotorasib or vehicle control (*n* ≥ 18 tumors/group). Unpaired *t* test was used in (B), (D), (E), (F), (L) and (M) to test for statistical significance: **** *p* < 0.0001; *** *p* < 0.001; ** *p* < 0.01;* *p* < 0.05. Error bars indicate SEM.

To extend our analysis beyond the KRAS(G12D) allele, we generated autochthonous *Kras^LSL-G12C/+^;Trp53^flox/flox^; Rosa26^tdTomato/+^* [*K(G12C)PT*] tumors and administered the clinically approved KRAS(G12C) allele-specific inhibitor sotorasib starting at 20 weeks PTI for 20 days (**Fig. 3K**). We observed a significant reduction in tumor burden (**Fig. 3L**). Similar to the KRAS(G12D) models, sotorasib promoted the enrichment of LUAD cells expressing the AT1 marker HOPX (**Fig. 3M**). Taken together, our results using orthogonal genetic and pharmacologic approaches indicate that alveolar differentiation is a prominent feature of LUAD cells that escape KRAS inhibition.

### AT1-like LUAD cells arise via trans-differentiation in response to KRAS inhibitors

We next sought to investigate the kinetics of the AT1-like LUAD cell state during KRAS inhibition. To do this, we introduced a “***MACD***” reporter construct into the *Hopx* locus of *KP* LUAD cells. *MACD* is a cDNA cassette comprising ***<u>m</u>****Scarlet* red fluorescent protein, a super-bright bioluminescence reporter [***<u>A</u>****kaLuc* (23)], tamoxifen-activatable Cre recombinase (***<u>C</u>****reER*), and diphtheria toxin (DT) receptor (***<u>D</u>****TR*) (**Fig. 4A; Supplementary Fig. 4A, B**). Into these cells, we also introduced a lentiviral construct encoding constitutive *Gaussia princeps* luciferase (G-Luc) and far-red monomeric iRFP670 (miRFP) fluorescent protein as well as a Cre-inducible tagBFP2 (BFP) fluorescent protein cassette. G-Luc is naturally secreted, enabling its detection using a specific substrate from a small volume of serum (24). Administration of MRTX1133 rapidly suppressed growth and G-Luc in the serum of mice bearing subcutaneous transplants, which resumed following cessation of MRTX1133 therapy (**Fig. 4B, C; Supplementary Fig. 4C**). We examined the kinetics of AT1-like state emergence in this experiment by AkaLuc bioluminescence imaging (**Fig. 4D**). Normalizing AkaLuc to G-Luc revealed a robust relative increase in *Hopx* gene expression in response to MRTX1133 (**Fig. 4E**), consistent with enrichment of AT1-like LUAD cells in residual disease. Interestingly, cessation of MRTX1133 administration led to a precipitous drop in relative *Hopx* expression (**Fig. 4E**), suggesting loss of AT1 identity (see below).

**Figure 4.**
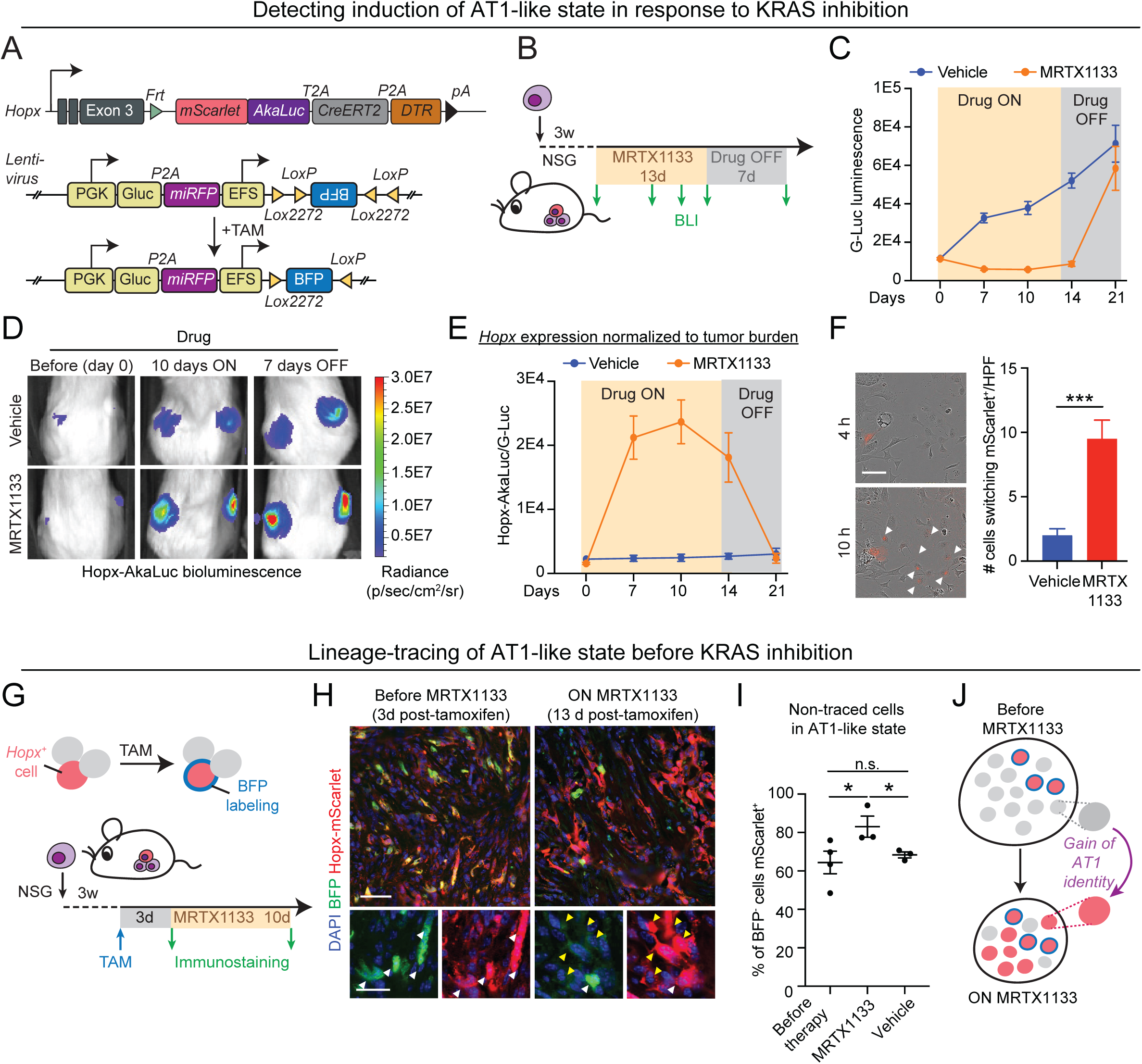
Enrichment of an AT1-like cancer cell state in response to KRAS inhibitors in LUAD. (**A**) Genetically engineered *Hopx-MACD* reporter system enabling lineage tracing and ablation of AT1-like *KP* LUAD cells. *Frt-Stop-Frt-mScarlet-Akaluc-CreERT2-DTR* reporter construct knocked in frame into the stop codon of *Hopx* exon 3. T2A and P2A: short polypeptide cleavage sites. *PGK-Gluc-miRFP-EFS-lox-BFP-lox* lentiviral lineage tracing vector was integrated into the *Hopx-MACD* reporter cells, enabling the lineage tracing of *Hopx^+^* cells with a single pulse of tamoxifen (TAM). (**B**) Outline of experimental design to investigate *Hopx* expression on and post KRAS(G12D) inhibition in subcutaneous *KRAS(G12D);P;Hopx-MACD* LUAD tumors using bioluminescence imaging (BLI) at the indicated time points. (**C**) Tumor burden measured by *Gaussia princeps* (G-Luc) luminescence in response to MRTX1133 therapy (*n* ≥ 7 mice/group). (**D**) Hopx-Akaluc bioluminescence detection in mice bearing *KP; Hopx-MACD* reporter allografts subjected to vehicle (*top*) or MRTX1133 (*bottom*) before (day 0), ON (10 days on), and OFF (7 days off) treatment. (**E**) Quantification of Hopx-Akaluc bioluminescence normalized to tumor burden (G-Luc bioluminescence) (*n* ≥ 14 tumors/group). (**F**) *Left*: Representative images of *KP; Hopx-MACD* reporter cells in 2D culture at 4 h and 10 h following exposure to MRTX1133. Note induction of *Hopx* expression as indicated by mScarlet fluorescence (white arrowheads). Scale bar: 200 µm. *Right*: Quantification of number of cells switching to the AT1-like state (mScarlet^+^) per high-power field (HPF) (n=6/group). (**G**) Experimental design for lineage-tracing the AT-like state before KRAS(G12D) inhibition in subcutaneous *KP; Hopx-MACD* reporter allografts. (**H**) Representative images of BFP and mScarlet immunofluorescence in the *KP; Hopx-MACD* reporter allografts before and on MRTX1133 or vehicle exposure. Scale bars: top 50 µm, bottom 25 µm. Note efficient labeling of mScarlet^+^ cells (BFP^+^/mScarlet^+^, white arrowheads) at 3 days following TAM administration and an increase of non-traced cells (BFP^-^/mScarlet^+^; yellow arrowheads) during MRTX1133 treatment. (**I**) Quantification of non-traced *Hopx^+^*(BFP^-^/mScarlet^+^) cancer cells in *Hopx^+^* (mScarlet^+^) tumors in (H) (*n* ≥ 3 tumors/group). (**J**) Schematic summary of findings: KRAS inhibition induces AT1 differentiation in non-AT1 LUAD cell states. Unpaired *t* test was used in (F) and (I) to test for statistical significance: *** *p* < 0.001; * *p* < 0.05. Error bars indicate SEM.

We next utilized our reporter system to interrogate whether the increase in AT1-like LUAD cells following KRAS inhibition is due to enrichment of pre-existing AT1-like cells or trans-differentiation of other LUAD cell states. We observed an induction of mScarlet fluorescence in cultured *KP; Hopx-MACD* reporter cells subjected to MRTX1133 (**Supplementary Fig. 4D**). Time-lapse imaging of individual cells revealed a significant increase in cells that switched from non-AT1-like (Hopx-mScarlet^-^) to AT1-like (mScarlet^+^) in response to MRTX1133 when compared to control (**Fig. 4F**). This indicates that KRAS inhibition induces *Hopx* in *Hopx^-^* LUAD cells. We next addressed the source of the AT1-like LUAD cells *in vivo* by lineage-tracing the AT1-like cells in established subcutaneous *KP; Hopx-MACD* allografts before administration of MRTX1133 (**Fig. 4G**). We observed faithful labeling of the AT1-like cells at 3 days following administration of a single pulse of tamoxifen (**Fig. 4H, left; Supplementary Fig. 4E**). Interestingly, the fraction of non-traced cells that express *Hopx* increased in response to MRTX1133 therapy, but not in response to vehicle (**Fig. 4H, I; Supplementary Fig. 4F**). Taken together, we conclude that KRAS inhibition induces AT1 trans-differentiation in LUAD cells (**Fig. 4J**).

### LUAD cells drive relapse but lose AT1-like identity following withdrawal of KRAS inhibition

Our findings raised the possibility that the AT1-like cells within the residual disease may serve as a source of relapse upon the emergence of acquired resistance. To model this scenario, we tested the growth potential of podoplanin^+^ AT1-like cells isolated from autochthonous *KPT* LUAD tumors at 20 days of MRTX1133 exposure *in vitro* in a tumor sphere assay (**Fig. 5A, B; Supplementary Fig. 5A**). MRTX1133 was withdrawn in these assays following cell sorting to mimic loss of drug activity on the mutant oncoprotein. We found that the podoplanin^+^ cells exhibited superior growth potential when compared to podoplanin^-^ cells isolated from the MRTX1133-treated tumors (**Fig. 5B, C**). No difference was observed when comparing podoplanin^+^ vs. podoplanin^-^ LUAD cells isolated from vehicle control tumors (**Fig. 5B, C**).

**Figure 5.**
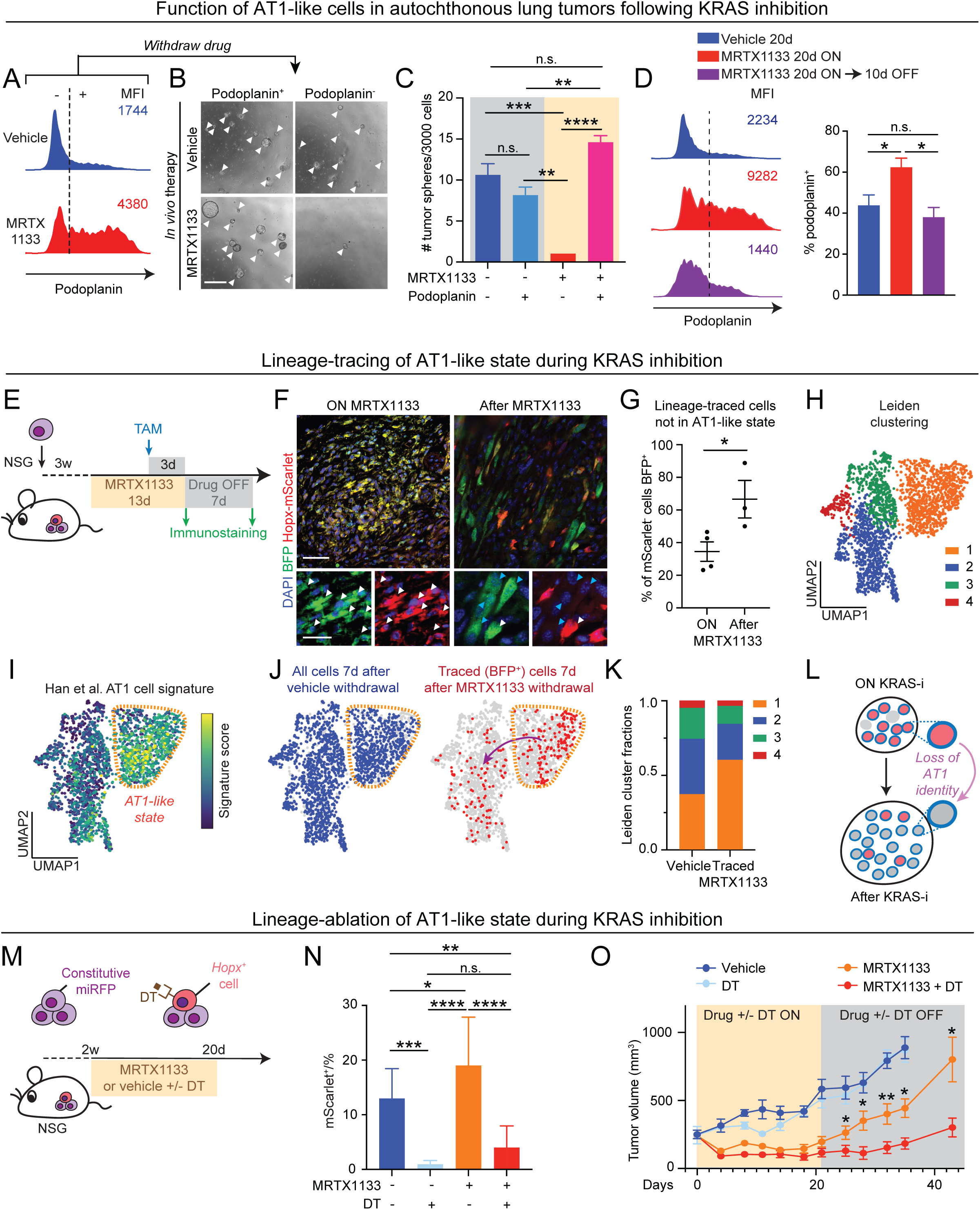
AT1-like LUAD cells drive resistance and relapse to KRAS inhibition. (**A**) Histogram showing podoplanin expression in tdTomato^+^/CD45^-^/CD31^-^/CD11b^-^/F480^-^/TER119^-^/DAPI^-^ (live) LUAD cells isolated from autochthonous tumors at 16 weeks post-tumor initiation plus 20 days on MRTX1133 (red) or vehicle control (blue). MFI: median fluorescence intensity. Dashed line separates podoplanin^-^ (-) and podoplanin^+^ (+) cells. (**B**) Representative images of 3D tumor spheres established from podoplanin^+^ [+ in (A)] and podoplanin^-^ [- in (A)] LUAD cells isolated from autochthonous tumors at 16 weeks post-tumor initiation plus 20 days on MRTX1133 or vehicle control. Spheres were cultured for 10 days in the absence of drug. Scale bar: 500 µm. (**C**) Quantification of the number of tumor spheres in the experiment outlined in (G-H); *n* = 8, 12, 3, and 5 mice from left to right. (**D**) *Left*: Histogram showing podoplanin expression in tdTomato^+^/CD45^-^/CD31^-^/CD11b^-^/F480^-^/TER119^-^/DAPI^-^ (live) LUAD cells isolated from autochthonous tumors at 16 weeks post-tumor initiation plus 20 days on vehicle (blue) or MRTX1133 (red), or 20 days on MRTX1133 plus 10 days of MRTX1133 washout (purple). *Right*: Quantification of the percentage of podoplanin^+^ cells in the aforementioned conditions (n>=6). One-way ANOVA was used to examine statistical significance. (**E**) Outline of experimental design to lineage-trace *Hopx^+^* AT1-like LUAD cells on and after MRTX1133 therapy vs. vehicle control in subcutaneous *KP; Hopx-MACD* reporter allografts. (**F**) Representative images of BFP and mScarlet immunofluorescence in *KP; Hopx-MACD* reporter allografts at 13 days on MRTX1133 therapy (left) and 7 days after relapse (right). Note the efficient labeling of traced cells (BFP^+^/mScarlet^+^; white arrowheads) after TAM administration and increase of lineage-traced cells that are not in the AT1-like state (BFP^+^/mScarlet^-^; turquoise arrowheads) after withdrawal of MRTX1133. Scale bar: Top: 50 µm. Bottom: 25 µm. (**G**) Quantification of lineage-traced cells not in AT1-like state (BFP^+^/mScarlet^-^) (*n* ≥ 3 tumors/group). An unpaired *t* test was used to examine significance. (**H**) Unsupervised clustering of miRFP^+^/CD45^-^/CD31^-^/CD11b^-^/F480^-^/TER119^-^/DAPI^-^ single LUAD cell transcriptomes, colored and annotated based on unsupervised Leiden clustering. (**I**) Projection of mouse AT1 cell gene expression signature (47) onto the UMAP space. (**J**) LUAD cell transcriptomes in the UMAP space following exposure to vehicle control (blue) or AT1 lineage-traced cells after 13 days of MRTX1133 exposure followed by 7 days of drug washout (red). Purple arrow indicates trans-differentiation of the AT1-like cells from the AT1-like state (orange dashed line) into other LUAD cell states. (**K**) Fraction of vehicle vs. AT1-lineage-traced cells transiently exposed to MRTX1133 in the Leiden clusters (H) (*n* = 3 mice/group). (**L**) Schematic summary of key findings: Loss of AT1-like identity during KRAS reactivation and tumor relapse. (**M**) Outline of experimental design to test ablation of *Hopx^+^* AT1-like LUAD cells in the context of MRTX1133 therapy or vehicle control in a subcutaneous *KP; Hopx-MACD* reporter allografts. (**N**) Quantification showing the percentage of mScarlet^+^ cells within the miRFP^+^/CD45^-^/CD31^-^/CD11b^-^/F480^-^/TER119^-^/DAPI^-^ (live) total LUAD cell pool at 20 days following MRTX1133 therapy or vehicle with or without DT (*n* ≥ 6). (**O**) Volume of subcutaneous *KP; Hopx-MACD* reporter allografts subjected to the indicated therapies; *n* ≥ 4/group. Asterisks indicate statistical significance (unpaired *t* test p-value) between MRTX1133 vs. MRTX1133 + DT groups. Two-way ANOVA was used in (C) and (N) to test for statistical significance: **** p < 0.0001; *** p < 0.001; ** p < 0.01; * p < 0.05. Error bars indicate SEM.

We had observed that cessation of MRTX1133 therapy leads to rapid regrowth of subcutaneous LUAD allografts, which coincided with downregulation of *Hopx* (**Fig. 4C-E; Supplementary Fig. 4C**). Similarly, we found that the proportion of the AT1-like cells was reduced back to baseline in autochthonous *KP* LUAD tumors at 10 days following cessation of a 20-day course of MRTX1133 therapy (**Fig. 5D**). These results suggested that the AT1-like LUAD cells de-differentiate following KRAS reactivation. To test this, we returned to our lineage-tracing system (**Fig. 4A, Supplementary Fig. 4A, B**). This time, we induced lineage-tracing of the AT1-like cells at residual disease (at 10 days of MRTX1133 therapy) (**Fig. 5E**). Again, our system traced the AT1-like LUAD cells at high fidelity (**Fig. 5F, left; Fig. 5G**). Interestingly, the majority of the traced cells lost AT1 identity upon withdrawal of MRTX1133 (**Fig. 5F, right; Fig. 5G; Supplementary Fig. 5B**). To provide a more granular view of the fate of the AT1-like cells following reactivation of KRAS, we performed scRNA-seq on the AT1 lineage-traced LUAD cells 7 days following withdrawal of MRTX1133 (**Fig. 5H, I**). Vehicle-treated LUAD cell transcriptomes were used as a reference (**Fig. 5J, left**). Not, although the traced cells showed an enrichment for the AT1-like state, a significant fraction of the traced cells were detected throughout the phenotypic space defined by vehicle-treated control cells (**Fig. 5J, K**). Thus, the AT1-like cells harbor the capacity to differentiate into multiple other LUAD cell states. In sum, these findings indicate the AT1-like LUAD cells within the residual disease harbor potential to reignite tumor growth upon treatment cessation or following emergence of acquired resistance. Furthermore, the reactivation of KRAS and relapse outgrowth are characterized by plastic transitions of the AT1-like cells into other cancer cell states within the tumors (**Fig. 5L**).

### Targeting AT1-like LUAD cell state augments response to KRAS inhibitors

Our results suggest that the AT1-like LUAD cells contribute to treatment resistance to KRAS inhibitors, casting the AT1-like cell state as an attractive therapeutic target for cytoablative therapies. To test this, we utilized the *KP; Hopx-MACD* reporter to ablate the AT1-like cells in established tumors by providing the mice with diphtheria toxin (DT) (**Fig. 5M**). MRTX1133 promoted the expression of the reporter construct, indicating enrichment of AT1-like cancer cells, whereas administration of DT efficiently eradicated the reporter-positive cells in both MRTX1133 and vehicle-treated tumors (**Fig. 5N**). Notably, co-administration of DT with MRTX1133 robustly delayed tumor relapse when compared to MRTX1133 alone, whereas ablation of the AT1-like cells alone had no effect on tumor growth in the absence of MRTX1133 (**Fig. 5O; Supplementary Fig. 5C**). These results indicate that targeting the AT1-like LUAD cells augments response to KRAS inhibitors.

### AT1-like cell state is enriched in residual disease induced by KRAS inhibitors in human LUAD

To evaluate whether our findings in the mouse model translate to human LUAD, we utilized four PDX models harboring *KRAS(G12C)* mutations (**Fig. 6A; Supplementary Fig. 6**) (25). We targeted KRAS(G12C) in these tumors using sotorasib or adagrasib, clinically approved small molecule inhibitors that specifically target the KRAS(G12C) oncoprotein (26). All four PDX models showed growth inhibition in response to the KRAS(G12C) inhibitors (**Fig. 6B; Supplementary Fig. 7A**). We observed similar AT1 differentiation and marker expression in the PDXs following KRAS(G12C) inhibition as in the mouse models (**Fig. 6C-E; Supplementary Fig. 7B, C**). Finally, we obtained clinical samples of KRAS(G12C) mutant human LUAD patient tissue longitudinally harvested from accessible metastatic sites either pre-treatment or on sotorasib or adagrasib therapy (**Supplementary Fig. 6**). Strikingly, patients on sotorasib or adagrasib showed a dramatic induction of AT1 differentiation in LUAD tissues as evidenced by HOPX immunoreactivity, whereas pre-treatment samples showed negligible HOPX expression (**Fig. 6F, G**). Interestingly, this phenotype was independent of pERK status in the tumors (**Supplementary Fig. 7D, E**). In conclusion, both human and mouse LUAD tumors respond to KRAS inhibition by undergoing a phenotypic shift towards an AT1-like state, indicating conservation of the cellular programs governing resistance to KRAS inhibition across species and individual patients (**Fig. 6H**).

**Figure 6.**
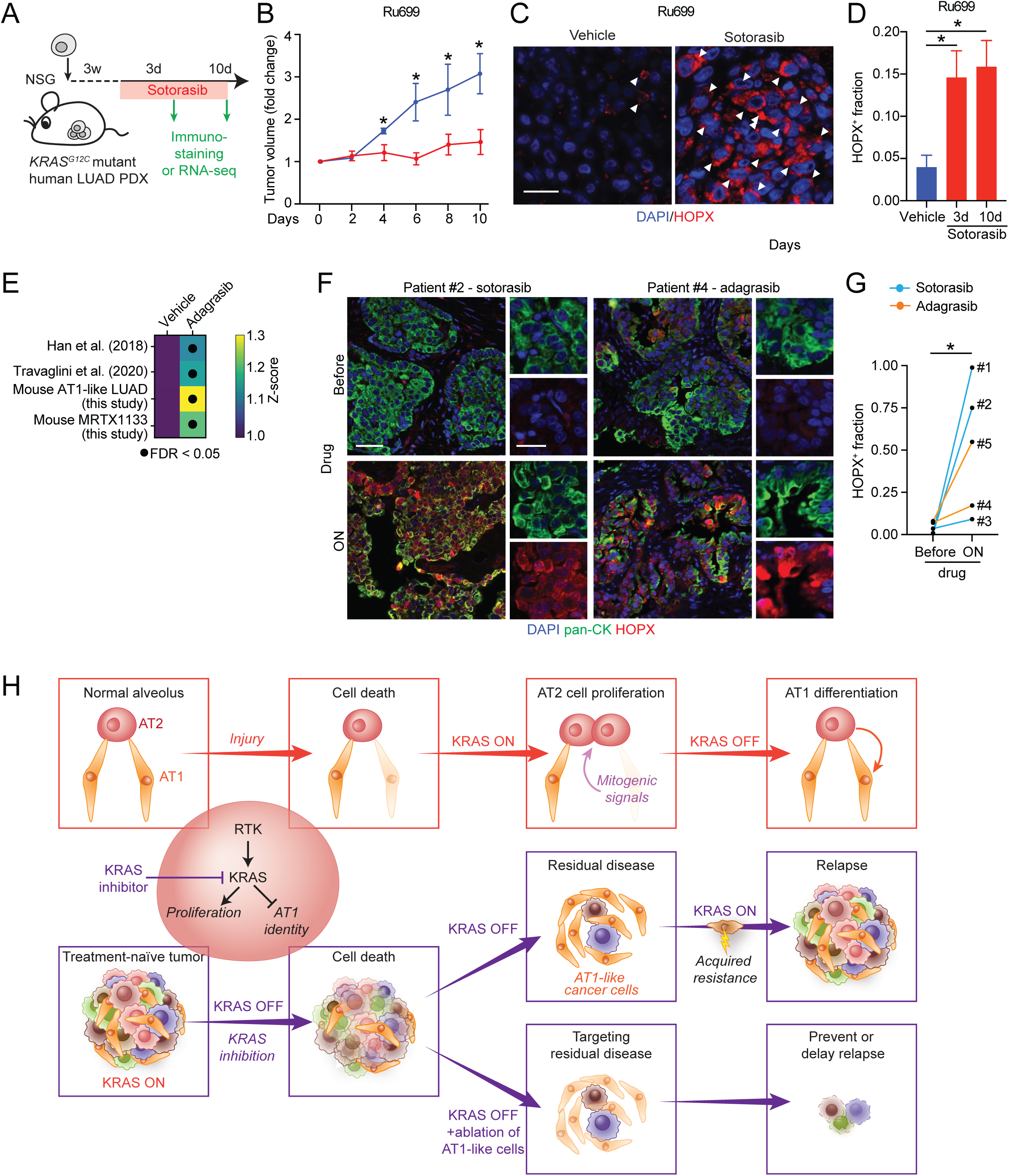
KRAS inhibitors enrich for AT1-like cancer cells in human LUAD. (**A**) Experiment to investigate KRAS(G12C) inhibition in a human LUAD PDX model using sotorasib. (**B**) Tumor volume fold change in response to sotorasib therapy (*n* ≥ 3 mice/group). Statistical significance was examined using an unpaired *t* test. (**C**) HOPX immunofluorescence in PDX models subjected to 10 days of sotorasib therapy. White arrowheads indicate HOPX^+^ cells. Scale bar: 20 µm. (**D**) Quantification of the fraction of HOPX^+^ cells in the PDX model (*n* = 6 tumors/experimental condition). An unpaired *t* test was used to test for statistical significance. (**E**) Signature scores of wild-type AT1 cell and AT1-like LUAD cell signatures in the PDXs following adagrasib therapy or vehicle control. “Mouse AT1-like LUAD” and “Mouse MRTX1133” signatures were defined in this study (**Supplementary Table 4** and **Supplementary Table 3**, respectively). Closed circle: *p* < 0.05 (three individual PDXs were used as biological replicates, see **Supplementary Fig. 6**). (**F**) HOPX and pan-cytokeratin (pan-CK) immunofluorescence in longitudinal tumor tissue biopsies obtained from human patients before and on sotorasib or adagrasib therapy. Scale bar: 50 µm. (**G**) Fraction of HOPX^+^/ pan-CK^+^ area in tumor regions in matched pre-treatment and on-sotorasib (blue line) or on-adagrasib (orange line) biopsies. A paired *t* test was used to test for statistical significance. (**H**) Schematic summary of key findings. See text. * p < 0.05. Error bars indicate SEM.

## Discussion

We discovered an AT1-like cancer cell state resistant to KRAS inhibition in LUAD. This finding is surprising because treatment-resistant cancer cell states have in many cancer types been suggested to harbor de-differentiated or stem-like features (10-13). Our previous work uncovered the enrichment of a de-differentiated high-plasticity cell state following chemotherapy in LUAD, implicating this cell state as a key driver of chemoresistance (9). Similarly, emerging data in human patients undergoing EGFR-or ALK-targeted therapy suggests that residual disease is enriched for a regenerative transcriptional signature indicative of a therapy-induced primitive cell state transition (14). Our results indicate residual disease induced by KRAS inhibitors is primarily composed of an AT1-like state that differentiates further towards the AT1 lineage with prolonged KRAS inhibition. Importantly, we observed similar accentuated AT1 differentiation upon silencing *Kras* during alveolar injury repair. In contrast, the primitive/de-differentiated cancer cell states are highly sensitive to KRAS inhibition, suggesting that KRAS signaling is an important driver of these states. Interestingly, the primitive regenerative state that emerges in human patients following EGFR or ALK inhibition also harbors some features of alveolar differentiation (14), suggesting that the mechanisms whereby KRAS suppresses alveolar differentiation may be at least partially shared with other oncogenic drivers. Identification of the signals and programs downstream of KRAS that control cancer cell fate is an important area of further inquiry and may elucidate molecular targets for suppressing mimicry of alveolar differentiation.

Our study uncovered a previously unknown function for KRAS in alveolar regeneration. Upon alveolar injury, mitogenic RTK ligands such as EGF and FGF7 are induced in the microenvironment of AT2 cells, promoting their proliferation (8). Our results cast KRAS as a central RAS species transducing these RTK signals. In parallel to the RTK ligands, the AT2 cells respond to proinflammatory cytokines such as interleukin-1β upon injury, which increases their plasticity and blocks AT1 differentiation (27). We found that suppressing *Kras* promotes AT1 differentiation of regenerative AT2 cells, suggesting that KRAS signaling is a barrier to AT1 differentiation during the regenerative response. These results imply that acquisition of plasticity during regeneration requires KRAS activity in addition to proinflammatory programs. Accordingly, wild-type KRAS may provide a therapeutic entry point in diseases that are characterized by chronic inflammation and incomplete alveolar repair, such as idiopathic pulmonary fibrosis or chronic obstructive pulmonary disease (8,27). Thus, our findings unexpectedly motivate repurposing of the emerging inhibitors of wild-type KRAS (1) for regenerative medicine beyond their intended use in oncology.

We observed that, in addition to the AT1-like state, the residual disease following KRAS(G12D) inhibition also contains AT2-like cancer cells. Notably, this cell state was not enriched following *shRNA*-mediated targeting of *Kras*, raising the possibility the allele-specific inhibitors that, unlike the *shRNAs*, largely spare wild-type KRAS may have distinct effects from genetic perturbation. Full genetic deletion of oncogenic KRAS was recently reported to lead to complete LUAD regression and lack of residual disease (28). The difference in our system as opposed to complete genetic ablation of oncogenic KRAS can be explained by a need for low level of KRAS activity in the AT1-like cells or for a short period of adaptation to KRAS inhibition that does not manifest in complete genetic knockouts.

Notably, we found that isolated AT1-like cancer cells possess significantly higher growth potential than the remaining pool of LUAD cells in residual disease, which is primarily composed of AT2-like cancer cells. Furthermore, lineage-tracing revealed that the AT1-like LUAD cells in residual disease are plastic, harboring the capacity to trans-differentiate into other LUAD cell states. These findings suggest that the AT1-like state may serve as a safe harbor for LUAD cells under therapeutic pressure until they evolve acquired resistance. One intriguing direction would be to investigate whether the plasticity of the AT1-like state endows it with the capacity for the neuroendocrine and/or squamous transformations that have been observed in response to EGFR and ALK-targeted therapies in LUAD (29). Another consideration is the susceptibility of the AT1-like state to immunological killing. KRAS has been shown to drive immunosuppression in lung cancer (30), whereas cytotoxic T lymphocytes contribute significantly to efficacy of KRAS-targeted therapies in pancreas cancer models (31). Additional studies are needed to investigate the combination of AT1 state ablation and KRAS inhibitors in immunocompetent models, either alone or in combination with immunotherapies.

Our data suggest that loss of KRAS signaling in LUAD cells repurposes a physiologic function – AT1 differentiation – in injury repair for drug resistance. Our reporter imaging and lineage-tracing results suggest that a significant proportion of the AT1-like cells observed in the residual disease arise from LUAD cells that are not in the AT1-like state, although the pre-existing AT1- like cells likely contribute as well. Such differentiation mimicry may have broader significance across other cancer types and oncoprotein-targeted therapies. Indeed, mutational activation of the mitogen-activated protein kinase (MAPK) pathway has been shown to drive de-differentiation in papillary thyroid cancer, which leads to downregulation of the iodine uptake machinery and resistance to radioiodine therapy (32,33). MEK inhibitors have limited efficacy as single agents in this disease, but they restore iodine import and re-sensitize the cancer cells to radioiodine in tumors harboring *BRAF* or *NRAS* mutations (32,33). Our data provide a conceptual framework for combining an oncoprotein-targeted therapy with a cytoablative therapy targeting a defined cancer cell state. Our work encourages the development of therapeutic strategies specifically targeting the AT1-like LUAD cells, such as CAR-T cells, antibody-drug conjugates, or engagers linking target cells with cytotoxic immune cells.

Taken together, our findings uncover an unexpected alveolar differentiation program underpinning resistance to KRAS inhibition in lung adenocarcinoma. We link this program to a previously unknown physiologic function for KRAS in lung regeneration. Our results implicate the AT1-like state as a biomarker of resistance to KRAS inhibitors and encourage development of therapeutic strategies to co-target KRAS and the AT1-like cancer cells in lung adenocarcinoma.

## Methods

### Mice

All animal studies were approved by the MSKCC Institutional Animal Care and Use Committee (protocol # 17-11-008). All genetically engineered mice were maintained on a mixed C57/BL6 and Sv129 background. Previously published *Kras^lox-stop-lox(LSL)-G12D/+^* (34)*, Trp53^flox/flox^* (35), Rosa26*^LSL-rtTA3-IRES-mKate2/+^* (36), Rosa26*^LSL-tdTomato/+^* (37), and Kras*^LSL-G12C/+^* (38) mouse strains were used in this study. In addition, three distinct mouse embryonic stem cell (mESC) lines harboring *TET-GFPshRNA* cassettes containing *shKras247*, *shKras462*, or *shRenilla* elements in the *Col1a1* locus were generated by recombinase-mediated cassette exchange, as previously reported (17,18,20). Sequences of the shRNAs are listed below:

- *shRenilla*: CAGGAATTATAATGCTTATCT
- *shKras247*: ACTTGACGATACAGCTAATTC
- *shKras462*: CTTGAAGATATTCACCATTA

Chimeric F0 mice were obtained by injecting donor ESCs into host embryos in the 8-cell stage and crossed into the Kras*^lox-stop-lox(LSL)-G12D/+^*; Trp53*^flox/flox^*; Rosa26*^LSL-rtTA3-IRES-mKate2/+^* background to generate Kras*^lox-stop-lox(LSL)-G12D/+^*; Trp53*^flox/flox^*; Rosa26*^LSL-rtTA3-IRES-mKate2/LSL-rtTA3-IRES-mKate2^*; *Col1a1^TRE-GFPshRNA/TRE-GFPshRNA^* (*KP; RIK; TET-GFPshRNA*) mice for experiments. *RIK; TET-GFPshRNA* mice without the *KP* alleles were used for the lung injury experiments. Mice were genotyped at ∼2 weeks of age. Primers are listed below:

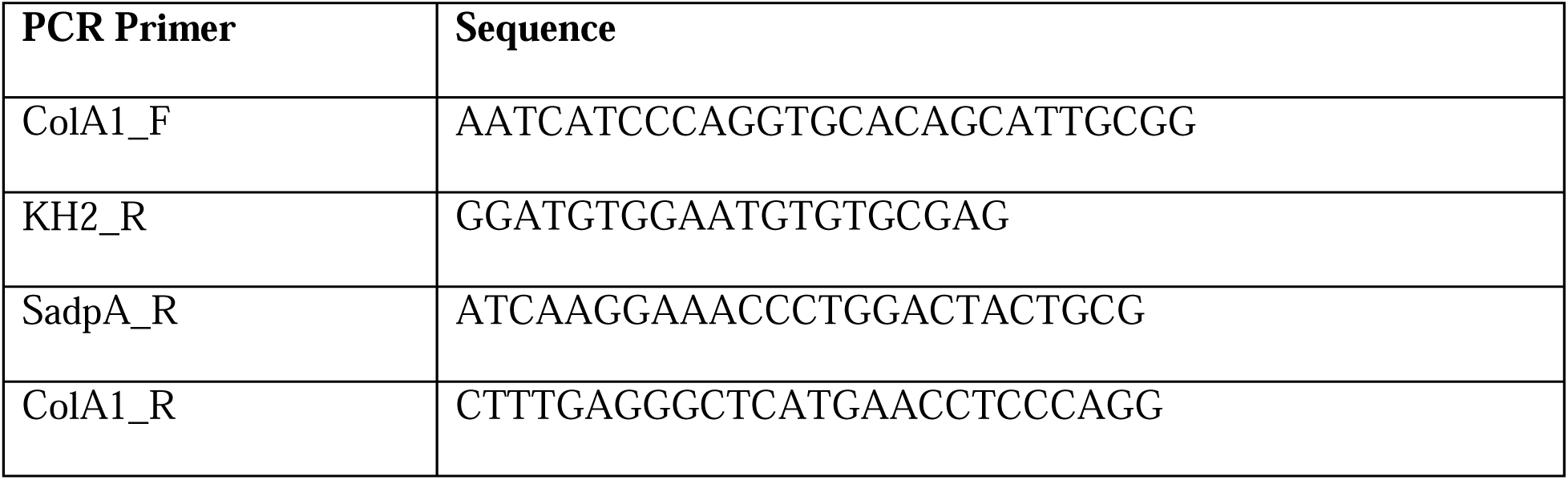

Immunocompromised NOD-SCID-gamma (NSG) mice were used as recipients for *KP* cell line allografts and LUAD PDXs. Mice were euthanized by CO_2_ asphyxiation followed by intracardiac perfusion with PBS to clear tissues of blood.

### Autochthonous lung adenocarcinoma models

Lung tumors were initiated in 8–12-week-old *KP; RIK; TET-GFPshRNA* and *KPT* mice by intratracheal administration of 2.5 x 10^8^ plaque-forming units (pfu) of AdSPC-Cre (#Berns-1168, University of Iowa) as previously described (39). Expression of GFP-linked *shRNAs* was induced at 16 weeks post-tumor initiation, a time point when a significant majority of the cancer cells had progressed from adenoma to adenocarcinoma, by placing the *KP; RIK; TET-GFPshRNA* mice on doxycycline hyclate chow (doxycycline 625 mg/kg, Envigo). To boost the expression of the *shRNA* constructs the mice were also provided with 25 mg/kg doxycycline hydrochloride (#D3447, Sigma-Aldrich) resuspended in PBS via intraperitoneal injection once a day for the first two days of doxycycline treatment.

### Patient-derived xenograft (PDX) models

PDXs were established under MSKCC IRB #06-107 and IRB #12-245. MSK-IMPACT focused exome profiling was performed on clinical samples as a part of routine care at MSKCC (40). Mutational analysis of all PDXs is provided in **Supplementary Fig. 6**. PDXs were dissociated into single cells with digestion buffer containing collagenase IV (#17104019, ThermoFisher Scientific, 0.1 U/ml), dispase (#354235, Corning, 0.6 U/ml), and DNase I (#69182–3; Sigma Aldrich, 10 U/ml) for 2 hours at 37 °C. Following enzymatic dissociation, samples were washed with 2% heat-inactivated FBS in S-MEM (#11380037, Thermo Fisher Scientific), filtered through a 100 μm cell strainer (#431752, Corning), and centrifuged at 1500 rpm for 5 min at 4 °C. The supernatant was removed, and the pellet was resuspended in lysis buffer (#555899, BD Biosciences) to remove red blood cells. Cells were passed through a 40 μm strainer (#431750, Corning) and centrifuged at 1500 rpm for 5 min at room temperature. Cells were resuspended in 90% of heat-inactivated FBS with 10% DMSO and frozen in liquid nitrogen. Before transplantation, PDXs were thawed and resuspended in S-MEM and mixed with Matrigel at a 1:1 ratio. 100,000 cells were implanted subcutaneously into both flanks of NSG mice.

### Administration of KRAS inhibitors

*KPT* mice bearing autochthonous LUAD tumors or NSG mice bearing subcutaneous *KP* cell line allografts were intraperitoneally administered freshly prepared MRTX1133 in captisol (#HY- 17031, MedChem Express) at 30 mg/kg BID, as previously described (7), starting at 16 weeks following initiation of autochthonous lung tumors or two weeks post-transplantation of cell lines. In addition, the NSG mice bearing *KP;Hopx-MACD* cell line allografts were administered 50 µg/kg diphtheria toxin (#D0564, Sigma) resuspended in sterile PBS every other day by intraperitoneal injection at three weeks post-transplantation. NSG mice bearing subcutaneous PDXs and *K(G12C)PT* autochthonous mice were administered sotorasib (#HY-114277, MedChem Express, 30 mg/kg QD) or adagrasib (100 mg/kg QD) by oral gavage for 3, 10 or 20 days, as previously described (26,41). Vehicle controls were used for all drug treatments.

### Hyperoxia lung injury

Alveolar injury was induced in *RIK; TET-GFPshRNA* mice by 66 hours of exposure to >85% O_2_ in a hyperoxia chamber (BioSpherix). The fractional-inspired oxygen concentration in the chamber was monitored by an in-line oxygen analyzer and maintained with a constant flow of gas (∼3 L/min). Following hyperoxia exposure, mice were placed on doxycycline chow for 7 days, followed by euthanasia and lung tissue harvest for histological analysis or FACS-based staining and isolation of lung epithelial cells for scRNA-seq (see below).

### Isolation of primary LUAD and lung epithelial cells

To dissociate tumors into single-cell suspensions, primary tumors were finely chopped with scissors and incubated with digestion buffer containing collagenase IV (#17104019, ThermoFisher Scientific, 0.1 U/ml), dispase (#354235, Corning, 0.6 U/ml), and DNase I (#69182–3; Sigma Aldrich, 10 U/ml). For dissociating normal lung epithelial cells, digestion buffer was administrated intratracheally into the lung as described before (27). Following enzymatic dissociation, samples were washed with 2% heat-inactivated FBS in S-MEM (#11380037, Thermo Fisher Scientific), filtered through a 100 μm cell strainer (#431752, Corning), and centrifuged at 1500 rpm for 5 min at 4 °C. The supernatant was removed, and the pellet was resuspended in lysis buffer (#555899, BD Biosciences) to remove red blood cells. Cells were passed through a 40 μm strainer (#431750, Corning) and centrifuged at 1500 rpm for 5 min at room temperature. Cells were resuspended in 2% of heat-inactivated FBS in PBS. Cell suspensions were blocked for 5 min at 4 °C with rat anti-mouse CD16/CD32 (#553142, Mouse BD Fc Block, BD Biosciences) in FACS buffer, and incubated for 30 min with a mix of four APC-conjugated antibodies binding CD45 (#103112, Biolegend), CD31 (#102410, Biolegend), CD11b (#101212, Biolegend), F4/80 (#123116, BioLegend), CD19 (#115512, Biolegend), and TER-119 (#116212, Biolegend). 300 nM DAPI was added as a live-cell marker. Individual cancer cell suspensions were incubated for 30 min with hashtag oligonucleotide-conjugated antibodies (see table below) in addition to FACS antibodies.

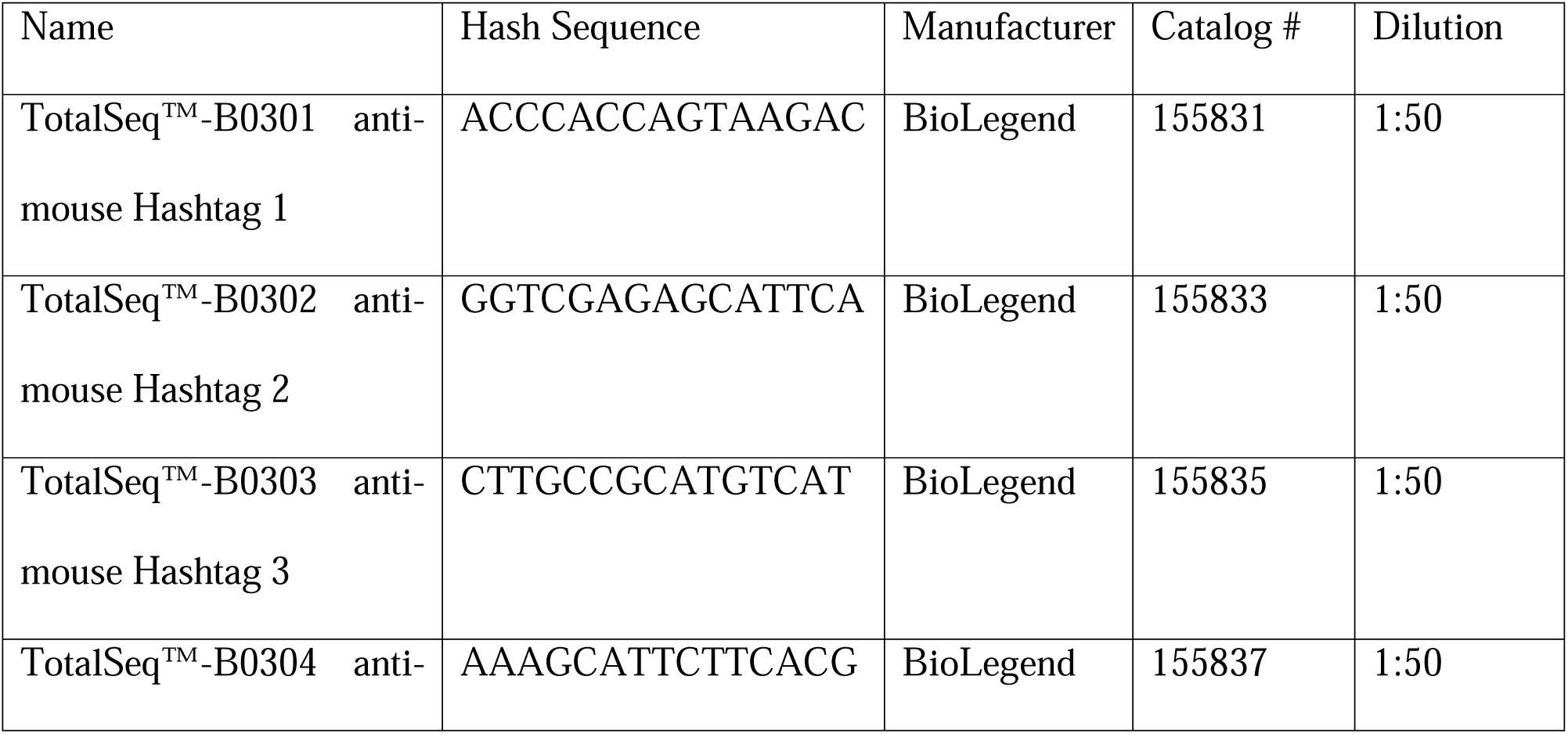
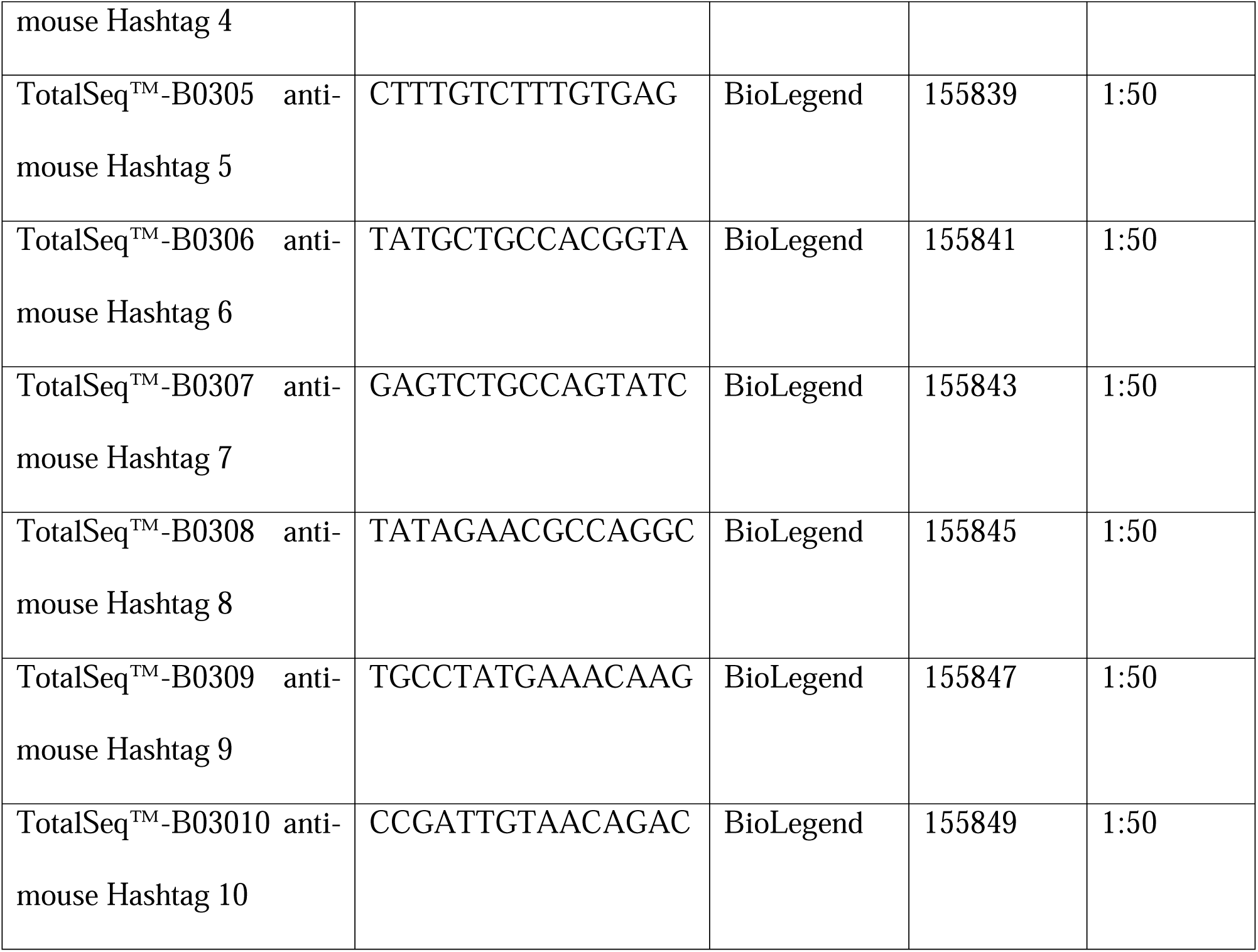

Cell sorting was performed on a FACSAria sorter (BD Biosciences). Sorted cells were collected directly into S-MEM with 2% of heat-inactivated FBS for scRNA-seq.

### Generation of Hopx reporter and lineage tracing cell line

Homology arms ∼1500 bp in length 5’ and 3’ to the end of *Hopx* exon 3 were amplified from C57/BL6 genomic DNA using high-fidelity PCR (KME-101, Toyobo). A homology-directed repair template donor vector where *frt-PGK-Hygro-pA-frt-T2A-mScarlet-Akaluc-T2A-CreER-P2A-DTR* is flanked by the homology arms was cloned into the pUC19 plasmid backbone (#638949, Takara). Donor vector and RNP complex were transfected into mESC line harboring *Kras^frt-stop-frt(LSL)-G12D/+^, Trp53^frt/frt^*by electroporation (4D Nucleofector, Lonza). Clones were validated by genotyping and sequencing using primers specifically detecting DTR, mScarlet, and spanning the homology arm. Primers are listed below:

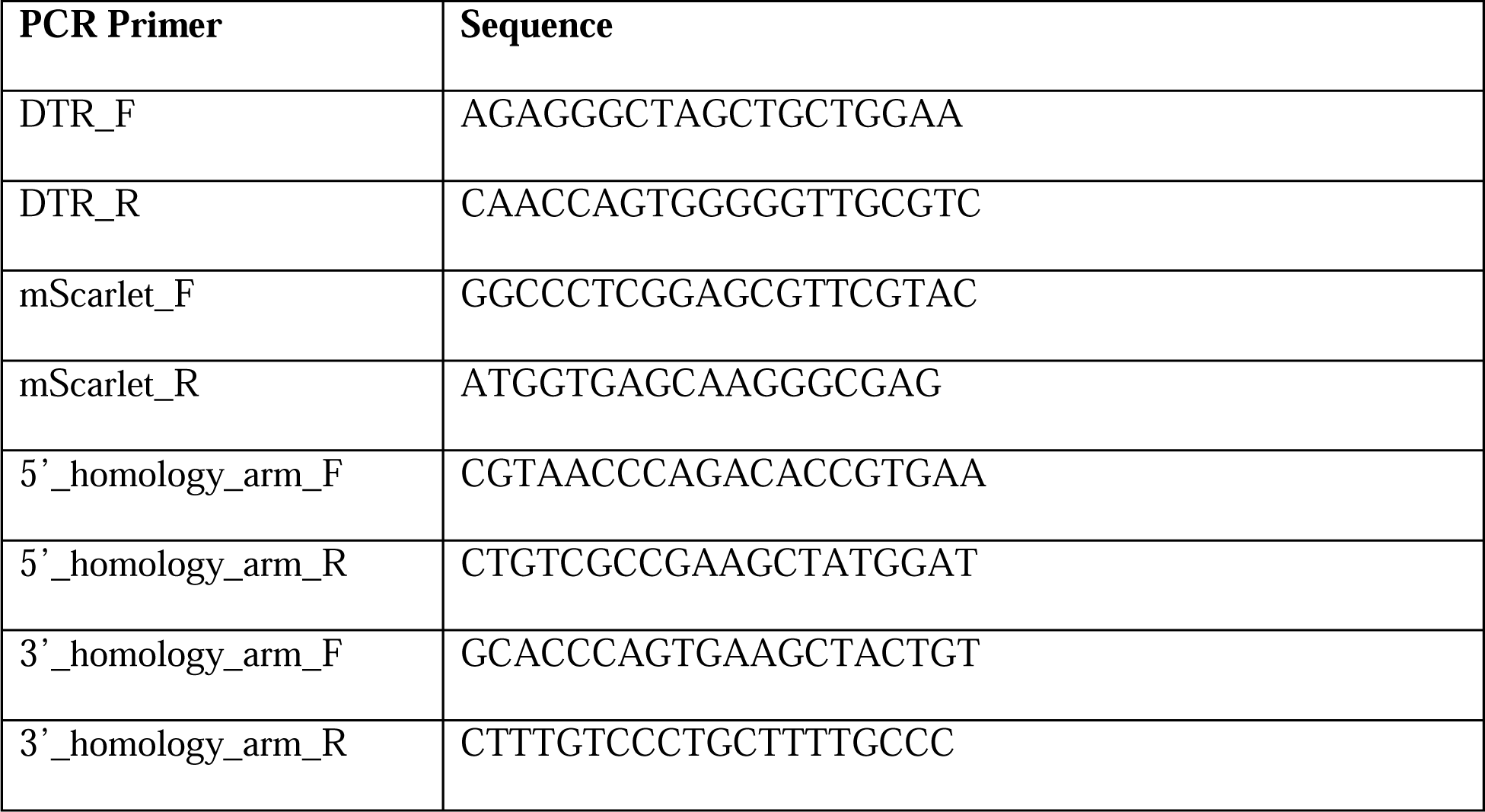

Chimeric F0 mice were obtained by injecting the donor ESCs into host embryos in the 8-cell stage. Mice were genotyped at ∼2 weeks of age (**Supplementary Fig. 4A,B**). Primers are listed below:

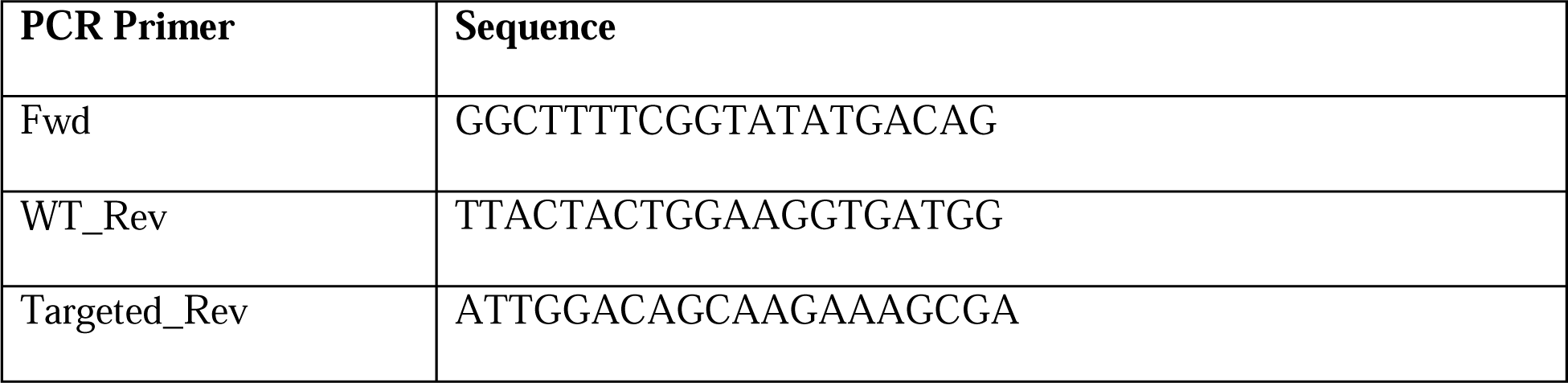

AT2 cells were isolated from F0 *Kras^frt-stop-frt(LSL)-G12D/+^; Trp53^frt/frt^; Hopx^MACD^*^/+^ – wild wild-type hybrids as described above. After sorting, the cells were transduced by lentivirus *PGK-Flpo* to remove the *frt-TAG-bGlobinpA-(PGK-Hygro-pA)i-frt* “STOP” cassette, and then the cells were cultured and expanded in tumor organoids (see below). Excision of the cassette was validated by genotyping using primers specifically detecting mScarlet-left homology arm and flow cytometry analysis detecting the presence of mScarlet fluorescence in a subset of the cells (**Supplementary Fig. 4A,B**).

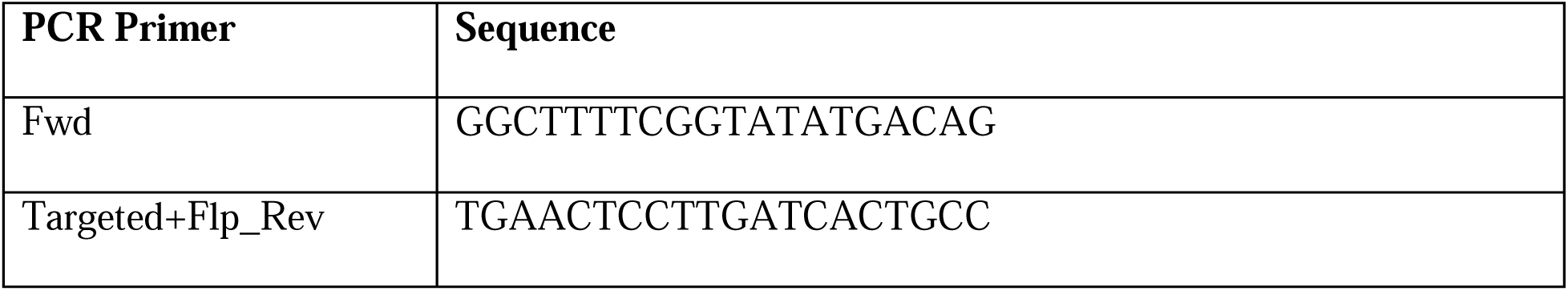

Lentiviral lineage tracing vector *PGK-Gluc-miRFP-EFS-lox-BFP-lox* (**Fig. 4A**) was integrated into the *Hopx-MACD* reporter tumor organoids by spin infection at 300 *g* for 30 min at 37C. Transduced cells were expanded and FACS-sorted into single-cell clones based on miRFP fluorescent expression, followed by expansion in culture.

### Cell culture and cell viability assays

Cells were cultured in a humidified incubator at 37°C under 5% CO_2_. Primary 2D mouse cell lines were generated from autochthonous *KP* and *KP; RIK; TET-GFPshRNA* LUAD tumors by dissociating primary lung tumors, as described above, and cultured in RPMI (Gibco) supplemented with 1% GlutaMax (#35050061, Thermo Fisher Scientific), 1% Pen/Strep (#15070063, Thermo Fisher Scientific) and 10% Heat-Inactivated FBS (#SH30910.03, Hyclone). Cells were passaged using 0.25% Trypsin-EDTA (#25200114, Invitrogen). Cell viability in culture was measured by MTT (3-(4,5-dimethylthiazol-2-yl)-2,5-diphenyltetrazolium bromide) assay. Cells were exposed to Dox for 72 h and incubated with 0.25 mg/ml MTT for 4 hours at 37˚C. Absorbance at OD 590 nm was measured to evaluate cell viability. An Incucyte imager (Sartorius) was used to image cell plates in both phase and RFP channels for time-lapse imaging. Cells were scanned every two hours for 3 days starting at 2 h from the start of MRTX1133 exposure.

### Tumor sphere culture

FACS-purified podoplanin^+^ or podoplanin^-^ LUAD cells isolated from *KPT* mice were resuspended in tumor sphere culture medium [Advanced DMEM/F12 (Gibco), 2% FBS (#SH30910.03, Hyclone), 1% GlutaMax (#35050061, Thermo Fisher Scientific), 1% Pen/Strep (#15070063, Thermo Fisher Scientific)] and mixed with 50 µl Matrigel (#CB-40230C, Fisher Scientific). Cell-Matrigel mix was placed in 24-well plates (# 353047, Thermo Fisher Scientific) and incubated in cell culture incubator. Tumor sphere culture media (800 µl) was added and replaced every 3 days during culture. Tumor spheres were imaged with an EVOS M5000 microscope (ThermoFisher).

### Tissue histology and immunofluorescence imaging

Tumors were harvested at the experimental endpoints, fixed in 10% neutral buffered formalin (Richard-Allan Scientific), embedded in paraffin, and cut into 5 µm sections. Slides were heated at 60°C for 30 minutes, deparaffinized, rehydrated with an alcohol series and incubated in Tris-EDTA antigen retrieval buffer (#E1161, Sigma-Aldrich) for 20 minutes in a pressure cooker at 95°C. Alternatively, formalin-fixed samples were incubated in 30% sucrose for 8-12 hours and frozen in OCT (#4585, Fisher HealthCare). Cryoblocks were sectioned at 5 µm thickness. Sections were placed on SuperFrost microscope slides (#12-550-15, Fischer Scientific) and used for staining. Sections were washed in PBS, permeabilized in 0.3% PBS-Triton X-100 for 40 minutes and blocked in PBS/0.1% Triton X-100 containing 2% BSA and 5% donkey serum (#D9663, Sigma-Aldrich). Primary antibodies were incubated overnight at 4°C in the blocking buffer. The following primary antibodies were used: RFP (#6g6, ChromTech, 1:500), GFP (#ab5450, Abcam, 1:500), HOPX (#sc-398703, Santa Cruz Biotechnology, 1:50), RAGE (#sc-365154, Santa Cruz Biotechnology, 1:100), SPC (#AB3786, EMD Millipore, 1:2000), pERK (#4370, Cell Signaling Technologies), and Ki67 (#14-5698-82, Thermo Scientific, 1:100), Pan-cytokeratin (#M3515292, Agilent Technologies, 1:1000). After washes in PBS tissues were incubated with fluorophore-conjugated secondary antibodies (#A-11055, #A-10042, #A-31571, ThermoFisher, 1:500) for 1 hour at room temperature. After staining slides were counterstained with DAPI (#D9542, Sigma Aldrich, 5 μg/ml) for 10 minutes and coverslipped with Mowiol mounting reagent. Hematoxylin and eosin (H&E) staining was performed using standard protocols. Images were acquired using 20x or 40x objectives on a Zeiss Axio Imager Z2 and ZEN 2.3 software or Mirax Midi-Scanner (Carl Zeiss AG).

### Immunohistochemistry

Immunohistochemistry was performed on 5 μm FFPE sections using standard staining protocols. Briefly, sections were de-paraffinized and heat-induced antigen retrieval was performed by EDTA antigen retrieval buffer (#E1161, Sigma Aldrich). Sections were blocked by BLOXALL solution (#SP-6000-100, Vector laboratories) at room temperature for 30 minutes and incubated with primary antibody at 4 °C overnight. IgG controls (#02-6102, #02-6202 and #10400C, Thermo Fisher Scientific) from the corresponding species of primary antibody were used as negative controls. Signal development was performed by ImmPRESS Polymer Detection Kits (#MP-7401-50, Vector Laboratories) and ImmPACT DAB Substrate Kit, Peroxidase (HRP) (#SK-4105, Vector Laboratories) following the manufacturer’s protocol. The sections were counterstained with hematoxylin (#72404, Thermo Fischer Scientific) and mounted with coverslips. The following primary antibodies were used: pERK (#4370, Cell Signaling Technologies, 1:1000). Mounted slides were digitally scanned using a Mirax Midi-Scanner (Carl Zeiss AG). Image analysis was performed using Fiji software.

### Capillary Western immunoassay (Wes)

Low-input protein size separation and immunodetection were performed with the capillary-based automated Western blot Wes system (# 031-108, ProteinSimple). Briefly, sorted cells were lysed in RIPA buffer (#9806, Cell Signaling Technology) and loaded at 1 µg into the Wes instrument for analysis. Primary antibodies against HOPX (#sc-398703, Santa Cruz Biotechnology, 1:50), KRAS(G12D) (#14429S, Cell Signaling Technology, 1:500), and GAPDH (#2118L, Cell Signaling Technology, 1:500) were detected using the appropriate HRP-conjugated secondary antibodies according to the manufacturer’s instructions.

### Western blotting

Cultured cells were rinsed with pre-cooled PBS and lysed by RIPA Buffer (#BP-115DG, Boston BioProducts) at 4□°C for 15□minutes, followed by centrifugation at 17,000 *g* for 15Lminutes. The supernatants were collected, quantified, and denatured for western blot analysis. Primary antibodies against KRAS(G12D) (#14429S, Cell Signaling Technology, 1:1000), Phospho-p44/42 MAPK (Erk1/2) (#4370, Cell Signaling Technology, 1:1000), ERK (#9102S, Cell Signaling Technology, 1:1000), GAPDH (#2118L, Cell Signaling Technology, 1:500) were used in the study.

### AkaLuc bioluminescence imaging *in vivo*

NSG mice bearing subcutaneous transplants of Hopx-MACD reporter cells were dosed with 100 µl of 30 mM of AkaLumine-HCl substrate resuspended in PBS (#808350, Millipore Sigma, TokeOni) and imaged on an IVIS Lumina II (Perkin Elmer)(23).

### Plasma sampling and Gaussia princeps luciferase measurements

Whole venous blood was harvested by puncturing the submandibular vein, followed by collection of 100 µl blood into collection vials (#02-675-185, Fisher Scientific) (42). Plasma was separated by centrifugation at >8000 *g* for 10 minutes at 4°C and diluted 1:10 in PBS. 200 µM *Gaussia* luciferase substrate coelenterazine-h (#301, NanoLight) was added, and luminescence was immediately measured on a BioTek Cytation 1 (Agilent).

### Lineage-tracing of Hopx^+^ cells

Subcutaneous *KP* LUAD cell line allografts harboring the *Hopx-MACD* allele and Flex-tagBFP lineage tracing vector (**Fig. 4A**) were established in NSG mice and monitored, as above. Tumor-bearing mice were administered one dose of tamoxifen (200 mg/kg by oral gavage) 3-weeks post-transplantation. Subcutaneous tumors were harvested 3 or 10 days after tamoxifen administration. The baseline measurement at 3 days was chosen to account for conversion of tamoxifen to the active metabolite 4-hydroxytamoxifen (4-OHT), recombination of the Flex-tagBFP in lineage-traced cells, and elimination of residual 4-OHT. Tamoxifen was dissolved in corn oil at 20 mg/mL in 55C for 1 hour, as before (39).

### Analysis of LUAD patients undergoing sotorasib and adagrasib therapy

Tissues from patients undergoing sotorasib (#1-3) or adagrasib (#4, #5) therapy were obtained under MSKCC Institutional Review Board approval (IRB #22-329, IRB #19-408). Date and anatomic site of tissue harvest are indicated below:

| **Patient** | **Pre-treatment tissue** | **On-treatment tissue** |
| --- | --- | --- |
| #1 | 4/5/2021, R humerus reamings | 9/20/2021, R kidney biopsy |
| #2 | 10/19/2021, L pleural nodule biopsies | 7/14/2022, L occipital brain tumors |
| #3 | 7/13/2022: T12 biopsy | 1/12/2023 T8 epidural tumor |
| #4 | 9/14/2020; Right supraclavicular mass | 5/11/2021, Right supraclavicular mass |
| #5 | 2/10/20, Liver Biopsy | 9/16/2020, Right lung nodule |

### Droplet-based scRNA-seq

Sorted cell suspensions were prepared for scRNA-seq using the 3′ v3 10X Genomics Chromium platform according to manufacturer’s instructions. Briefly, sorted cells were washed once with PBS containing 1% bovine serum albumin (BSA) and resuspended in PBS containing 1% BSA to a final concentration of 700–1,300 cells per μl. The viability of cells in all experiments was above 80%, as confirmed with 0.2% (w/v) Trypan Blue staining (Countess II, Invitrogen). Cells were captured in droplets. Following reverse transcription and cell barcoding in droplets, emulsions were broken, and cDNA purified using Dynabeads MyOne SILANE followed by PCR amplification as per manufacturer’s instructions. Between 20,000 to 25,000 cells were targeted for each droplet formulation lane. Samples were multiplexed together in the lanes following the TotalSeq B cell hashing protocol (43). Final libraries were sequenced on Illumina NovaSeq S4 platform (R1 – 28 cycles, i7 – 8 cycles, R2 – 90 cycles).

#### Computational analyses

### Analysis of single-cell RNA-sequencing data

FASTQ files of single-cell RNA-sequencing data generated on the 10X Chromium platform were processed using the standard CellRanger pipeline (version 5.0.0). Reads were aligned to a custom GRCm38 / mm10 reference including the *tdTomato*, *GFP*, and *mKate* transgenes used in this study. Cell-gene count matrices were analyzed using a combination of published packages and custom scripts centered around the scanpy / AnnData ecosystem (44). Single-cell RNA-sequencing datasets from *GFPshRNA KP* LUAD tumors, MRTX1133-treated *KP* LUAD tumors, and lung regeneration experiments were analyzed separately using the same workflows.

Single-cell RNA-sequencing data from treated and control replicates were dehashed and compiled into a combined count matrix. Cells with less than 500 UMIs, more than 20% mitochondrial UMIs, and low complexity based on the number of detected genes vs. number of UMIs were removed. Doublets were filtered using scrublet (45). UMI counts were normalized using the size factor approach described by Lun et al. (46).

Highly variable features were selected using a variance stabilizing transformation and dimensionality reduction was performed on normalized, log2-transformed count data using principal component analysis. The dimensionality reduced count matrices were used as input for UMAP-embedding and unsupervised clustering with the leiden algorithm; *bbknn* was used to control for batch effects.

To ensure that treatment does not bias cell state assignment, unsupervised clustering was performed only on control cells (*shRenilla* or vehicle-treated) and used as input to train a logistic regression (logit) classifier which was used to assign each cell to a cell state. This approach ensured that the cell states defined in the study are representative of states previously identified in wildtype tumors (9).

Normalized expression data was first MaxAbs scaled to give each gene equal weight. Gene signature scores were then calculated using the *score_genes* function in *scanpy*. Normal adult AT1 cell signatures were compiled from Angelidis et al. (47), Han et al. (47), Strunz et al. (48), and Travaglini et al. (49). KRAS activity signature genes were taken from HALLMARK_KRAS_SIGNALING_UP.v2022.1.Mm (50).

### Bulk RNA sequencing analysis

PDXes and autochthonous *KP* LUAD tumors were FACS-isolated, RNA was extracted, and libraries were prepared for SMARTerSeq2, n = 3 mice/group. FASTQ files were processed using the standard DESeq2 pipeline (version 1.24.0). Reads were aligned to the GRCm38 mouse genome. Count matrices were generated was performed using DESeq2. Data were filtered to only include genes with 10 or more counts in 2 or more samples. The significance of differential expression between vehicle and treatment groups was quantified by one-way ANOVA.

## Data Availability

The data generated in this study are available within the article and its supplementary data files. All other raw data are available upon request from the corresponding author.

## Authors’ Contributions

**Z. Li:** Conceptualization, investigation, data curation, formal analysis, validation, methodology, visualization, writing–original draft, writing–review and editing. **X. Zhuang:** Conceptualization, investigation, data curation, formal analysis, validation, methodology, visualization, writing– review and editing. **C.-H. Pan:** investigation, data curation, formal analysis, validation, methodology, visualization, writing–review and editing. **Y. Yan:** Resources, investigation, validation, methodology. **R. Thummalapalli:** Resources, validation. **J. Hallin:** Resources, formal analysis, validation. **S. Torborg:** Data curation, formal analysis, resources, validation. **A. Singhal:** Resources, data curation. **J. C. Chang:** Resources, investigation. **E. Manchado:** Resources, investigation. **L. E. Dow:** Resources, methodology. **S. W. Lowe:** Resources, supervision, methodology. **R. Yaeger:** Resources, methodology. **J. G. Christensen:** Resources, supervision, methodology. **C. M. Rudin:** Resources, supervision, funding acquisition. **S. Joost:** Conceptualization, data curation, formal analysis, supervision, methodology, visualization, writing–review and editing. **T. Tammela:** Conceptualization, resources, data curation, formal analysis, supervision, funding acquisition, validation, methodology, visualization, writing– original draft, writing–review and editing.

## Acknowledgments

We thank J. Fagin, N. Rosen, and members of the Tammela laboratory for helpful discussions; Y.-C. Han for critical comments on the manuscript; E. de Stanchina and the Antitumor Assessment Core Facility for support with drug administration and tumor transplant experiments; members of the Molecular Diagnostics Service in the Department of Pathology for generating clinical MSK-IMPACT data; R. Gardner, K. Daniels, and A. Longhini for FACS support; K. Manova and M. Tipping for histology support; N. Mohibullah for next-generation sequencing; H. Alcorn and O. Grbovic-Huezo for laboratory management; and M. Blum, J. Chan, S. Ding,M. Gregory, G. Hartmann, A. Hudson, H. Styers, C. Sussman, and K. Wu for help with experiments; and S. Weil for artwork in Figure 6H. This work was supported by the American Cancer Society and a Josie Robertson Scholarship (to T.T.); the HHMI and NIH/NCI R01- CA233944 (to S.W.L.); and by the NIH/NCI Cancer Center Support Grant P30-CA08748 (to MSKCC). X. Z. received support from a training award from New York Stem Cell Science NYSTEM (#C32559GG) at the Center for Stem Cell Biology at MSKCC; S.J. from the Hope Funds for Cancer Research; and S.T. from NIH/NCI (F30-CA254120) and a Medical Scientist Training Program grant from the National Institute of General Medical Sciences of the National Institutes of Health under award number T32GM007739 to the Weill Cornell/Rockefeller/Sloan Kettering Tri-Institutional MD-PhD Program. We acknowledge the use of the Antitumor Assessment, Integrated Genomics Operation, Flow Cytometry, and Molecular Cytology Core Facilities at the Sloan Kettering Institute, funded by CCSG P30-CA08748.

## Legends to Supplementary Figures

***Supplementary Figure 1. Generation and validation of genetically engineered mouse model enabling inducible shRNA-based targeting of Kras in autochthonous LUAD tumors.*** (**A**) PCR genotyping of the *Col1a1* homing cassette (CHC) locus in mouse embryonic stem cells (mESCs) targeted with Renilla control or *Kras*-targeting *shRNAs*. CHCe: empty *Col1a1* homing cassette locus; CHCt: targeted *Col1a1* homing cassette locus; WT: wild-type. (**B**) Validation of knockdown efficiency of shRNAs targeting *Kras* in mESCs by Western blotting for KRAS following 96 hours of Dox exposure. (**C**) PCR genotyping of CHC locus in F1 *KP; RIK; TET-GFPshRNA* mice harboring control or *Kras*-targeting *shRNAs*. (**D**) Outline of cell line generation from autochthonous *KP; RIK; TET-GFPshRNA* LUAD tumors. (**E**) Immunoblotting for KRAS and pERK in cultured *KP; RIK; TET-GFPshRNA* LUAD cells exposed to Dox or vehicle control for 48 h. (**F**) Cell viability of cultured *KP; RIK; TET-GFPshRNA* LUAD cells by MTT (3-(4,5- dimethylthiazol-2-yl)-2,5-diphenyltetrazolium bromide) assay following 72 h exposure to Dox or vehicle. (**G**) Detection of KRAS(G12D) protein in sorted GFP^+^/mKate^+^/CD45^-^/CD31^-^/CD11b^-^/F480^-^/TER119^-^/DAPI^-^ (live) primary *KP; RIK; TET-GFPshRNA* LUAD cells by capillary immunoassay after 20 days of Dox exposure. (**H**) Representative Ki67 and GFP immunofluorescence in autochthonous *KP; RIK; TET-GFPshRenilla* and *KP; RIK; TET-GFPshKras247* LUAD tumors after 20 days of Dox exposure. Note loss of GFP expression and emergence of proliferating (Ki67^+^) cancer cells in the *shKras247* tumor, demarcated by a dashed line. Scale bar: 100 µm. (**I**) Quantification of apoptotic (cleaved caspase 3 (CC3)^+^/tdTomato^+^) cancer cells at 16 weeks post-tumor initiation + 3, 10, or 20 days on MRTX1133 or vehicle control (*n* ≥ 96, 32, and 34 tumors for each time point, respectively). One-way ANOVA was used in (F); An unpaired *t* test was used in (I) to test for statistical significance: **** *p* < 0.0001; *** *p* < 0.001; ** *p* < 0.01. Error bars indicate SEM.

***Supplementary Figure 2. KRAS signaling inversely correlates with AT1 marker expression.*** (**A**) *Left*: Signature score of two independent previously reported LUAD residual disease signatures in the AT1-like LUAD cell state (orange) in each model. The average score of the single-cell transcriptomes is shown in each group. Open circle: *p* < 0.05; closed circle: *p* < 0.01 (unpaired *t* test shKras vs. shRenilla with individual tumors as biological replicates). *Right*: Unsupervised clustering of GFP^+^/mKate^+^ single LUAD cell transcriptomes, colored and annotated based on Marjanovic et al. (9). Normal healthy AT2 and AT1 single-cell transcriptomes (gray) isolated from wild-type mice are co-embedded. (**B**) Detection of ∼8 kDa HOPX protein in sorted GFP^+^/mKate^+^/CD45^-^/CD31^-^/CD11b^-^/F480^-^/TER119^-^/DAPI^-^ (live) primary *KP; RIK; TET-GFPshRNA* LUAD cells by capillary immunoassay after 20 days of Dox exposure. (**C**) Quantification of HOPX protein level in (B). (**D**) *Kras* gene expression in the distinct cancer cell states in unperturbed autochthonous *KP* LUAD tumors at 16 weeks post-tumor initiation. (**E**) KRAS gene expression signature score in the distinct cancer cell states in unperturbed autochthonous *KP* LUAD tumors at 16 weeks post-tumor initiation. Gene set: HALLMARK_KRAS_SIGNALING_UP.v2022.1.Mm.

***Supplementary Figure 3. KRAS inhibition but not chemotherapy induces AT1 differentiation in LUAD.*** (**A**) Signature score of two independent residual disease signatures in the AT1-like LUAD cell state (orange) in *KP* LUAD tumors following vehicle or MRTX1133 administration. The average score of the single-cell transcriptomes is shown in each group (n=3 mice/group). Open circle: *p* < 0.05; closed circle: *p* < 0.01 (**B**) Histogram showing podoplanin expression in EpCAM^+^/CD45^-^/CD31^-^/CD11b^-^/F480^-^/TER119^-^/DAPI^-^ (live) LUAD cells isolated from autochthonous tumors at 16 weeks post-tumor initiation plus 20 days on cisplatin (red, 5mg/kg i.p. administration weekly) or vehicle control (blue). MFI: median fluorescence intensity. Dashed line separates podoplanin^-^ (-) and podoplanin^+^ (+) cells. (**C**) Quantification of the fraction of podoplanin^+^ cells in (B) (n>=3). (**D**) Signature scores of wild-type AT1 cell and AT1-like LUAD cell signatures in the autochthonous *KP* model following cisplatin therapy or vehicle control. “Mouse AT1-like LUAD” and “Mouse MRTX1133” signatures were defined in this study (**Supplementary Table 4** and **Supplementary Table 3**, respectively). No significant difference was detected.

***Supplementary Figure 4. Allele-specific inhibition of KRAS(G12D) promotes AT1 differentiation in mouse LUAD models.*** (**A**) *Frt-Stop-Frt-mScarlet-Akaluc-CreERT2-DTR* (*MACD*) reporter construct knocked into the stop codon of *Hopx* exon 3 in *Kras^frt-stop-frt-G12D/+^; Trp53^frt/frt^* mouse embryonic stem cells (mESCs). AT2 cells isolated from F0 hybrid mice were transduced *in vitro* with lentiviral codon-optimized Flp (PGK-FlpO) to remove the *frt-stop-frt* cassette, activate KRAS(G12D), and delete both copies of Trp53, transforming the cells and poising the *Hopx-MACD* reporter. T2A and P2A: short polypeptide cleavage sites. (**B**) PCR genotyping of the *Hopx-MACD* reporter in WT or targeted mESCs or FlpO-transduced *KP; Hopx-MACD* reporter cells. (**C**) Volume of subcutaneous *KP; Hopx-MACD* reporter allografts, subjected to the indicated therapies (n >= 14). (**D**) mScarlet^+^ fraction in 2D *KP; Hopx-MACD* reporter cell culture exposed to vehicle or MRTX1133 at indicated timepoints (*n* ≥ 8/group). (**E**) Histogram showing BFP expression in miRFP^+^/CD45^-^/CD31^-^/CD11b^-^/F480^-^/TER119^-^/DAPI^-^ (live) LUAD cells isolated from subcutaneous tumors at 3 weeks post-implantation with (+TAM) or without (-TAM) a single dose of tamoxifen. (**F**) Histogram showing BFP expression in mScarlet+/miRFP^+^/CD45^-^/CD31^-^/CD11b^-^/F480^-^/TER119^-^/DAPI^-^ (live) LUAD cells isolated from subcutaneous tumors at 3 weeks post-implantation plus 3 days post-tamoxifen (top) or tamoxifen plus 10 days of MRTX1133 exposure (13 days post-tamoxifen).

***Supplementary Figure 5. AT1-like LUAD cells drive resistance to KRAS inhibitors and tumor relapse following treatment cessation.*** (**A**) Projection of *Pdpn* expression onto the UMAP in (Fig. 3G). (**B**) Histogram showing mScarlet expression in BFP+/miRFP^+^/CD45^-^/CD31^-^/CD11b^-^/F480^-^/TER119^-^/DAPI^-^ (live) LUAD cells isolated from *KP; Hopx-MACD* reporter allografts at 3 weeks post-implantation plus 10 days on MRTX1133 treatment plus 3 days post tamoxifen induction or 7 days off MRTX1133 treatment. (**C**) Representative photograph of subcutaneous *KP; Hopx-MACD* reporter allografts at the conclusion of the experiment outlined in (**Fig. 5M**). Scale bar: 5 mm.

***Supplementary Figure 6. Mutational spectra of human PDX models and patient samples used in the study.***

***Supplementary Figure 7. Allele-specific oncogenic KRAS(G12C) inhibitors promote an AT1- like cancer cell state in human LUAD.*** (**A**) Growth kinetics of PDX models subjected to adagrasib therapy (n=5/group). (**B**) Representative images of RAGE immunofluorescence in PDX model Ru699 subjected to 10 days of sotorasib therapy. White arrowheads indicate RAGE^+^ cells. Scale bar: 20 µm. (**C**) Quantification of the fraction of RAGE^+^ cells in the PDX model (*n* = 6 tumors/experimental condition). (**D**) Fraction of pERK^+^/pan-CK^+^ area in tumor regions in matched pre-treatment and on-sotorasib (blue line) or on-adagrasib (orange line) biopsies. (**E**) Scatter plot showing the fraction of pERK^+^/pan-CK^+^ and HOPX^+^/pan-CK^+^ in matched pre-treatment and on-sotorasib (blue line) or on-adagrasib (orange line) biopsies. The arrows point from pre-treatment sample to on-treatment sample.

***Supplementary Table 1:*** Differential gene expression analysis of autochthonous *KP* LUAD cells expressing *shRen*illa or *shKras*.

***Supplementary Table 2:*** Differential gene expression analysis of lineage-traced AT2 cell progeny expressing *shRenilla* or *shKras* following hyperoxia injury.

***Supplementary Table 3:*** Differential gene expression analysis of autochthonous *KP* LUAD cells following treatment with vehicle or MRTX1133.

***Supplementary Table 4:*** Mouse AT1-like LUAD signature genes.

