## Supplementary Figures for "Alveolar differentiation drives resistance to KRAS inhibition in lung adenocarcinoma"

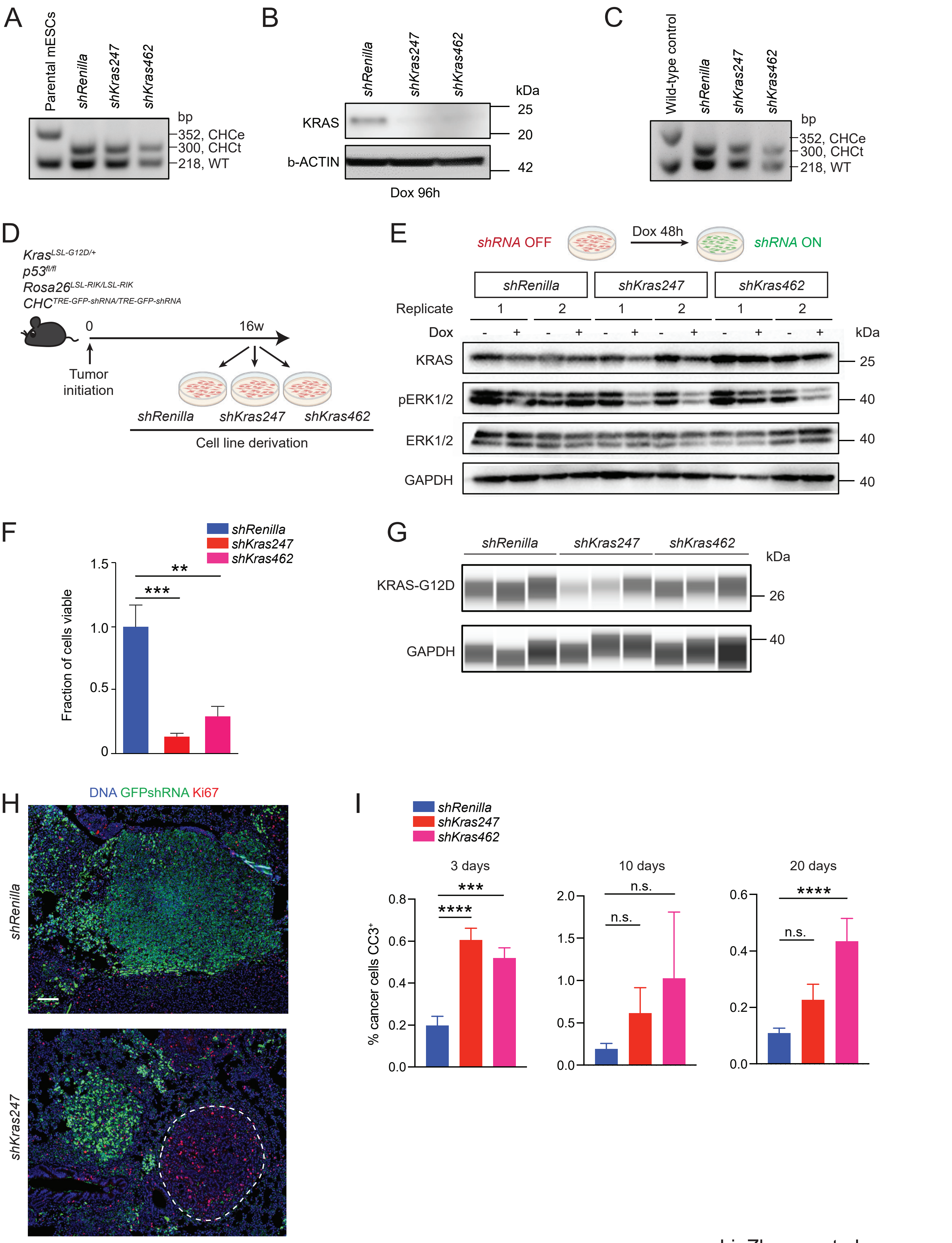
A
Parental mESCs
shRenilla
shKras247
shKras462
bp
352, CHCe
300, CHCt
218, WT
B
shRenilla
shKras247
shKras462
kDa
KRAS
b-ACTIN
25
20
42
Dox 96h
C
Wild-type control
shRenilla
shKras247
shKras462
bp
352, CHCe
300, CHCt
218, WT
D
Kras LSL-G12D/+
p53 fl/fl
Rosa26 LSL-RIK/LSL-RIK
CHC TRE-GFP-shRNA/TRE-GFP-shRNA
0
16w
Tumor initiation
shRenilla
shKras247
shKras462
Cell line derivation
E
shRNA OFF
Dox 48h
shRNA ON
shRenilla
shKras247
shKras462
Replicate
1
2
1
2
1
2
Dox
- + - + - + - + - + - +
kDa
KRAS
pERK1/2
ERK1/2
GAPDH
25
40
40
40
F
shRenilla
shKras247
shKras462
Fraction of cells viable
1.5
1.0
0.5
0
**
***
G
shRenilla
shKras247
shKras462
kDa
KRAS-G12D
GAPDH
26
40
H
DNA GFPshRNA Ki67
shRenilla
shKras247
I
shRenilla
shKras247
shKras462
3 days
10 days
20 days
% cancer cells CC3+
0.8
0.6
0.4
0.2
0.0
***
****
2.0
1.5
1.0
0.5
0.0
n.s.
n.s.
0.6
0.4
0.2
0.0
****
n.s.

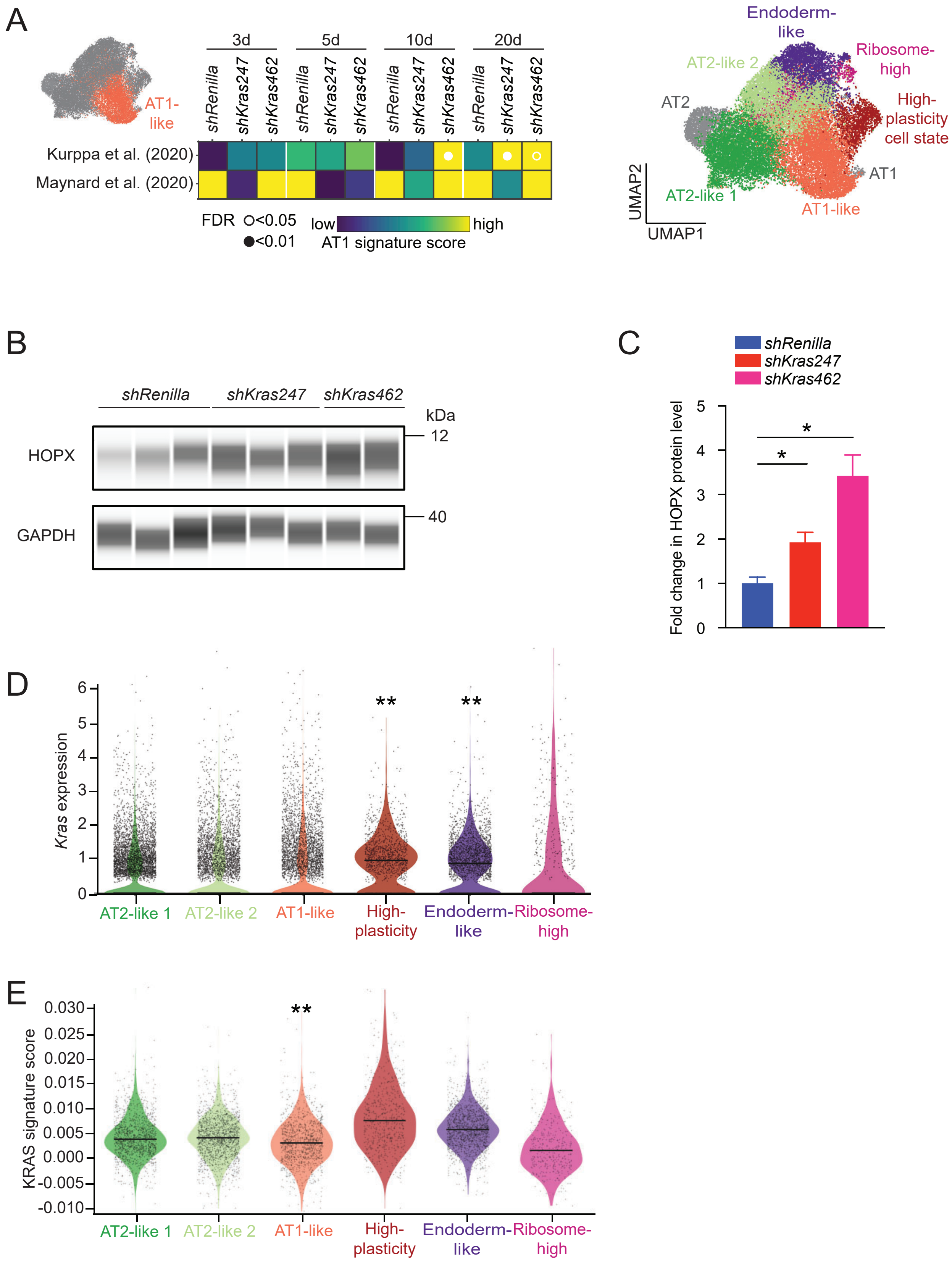


Li, Zhuang et al.
Supplementary Figure 2

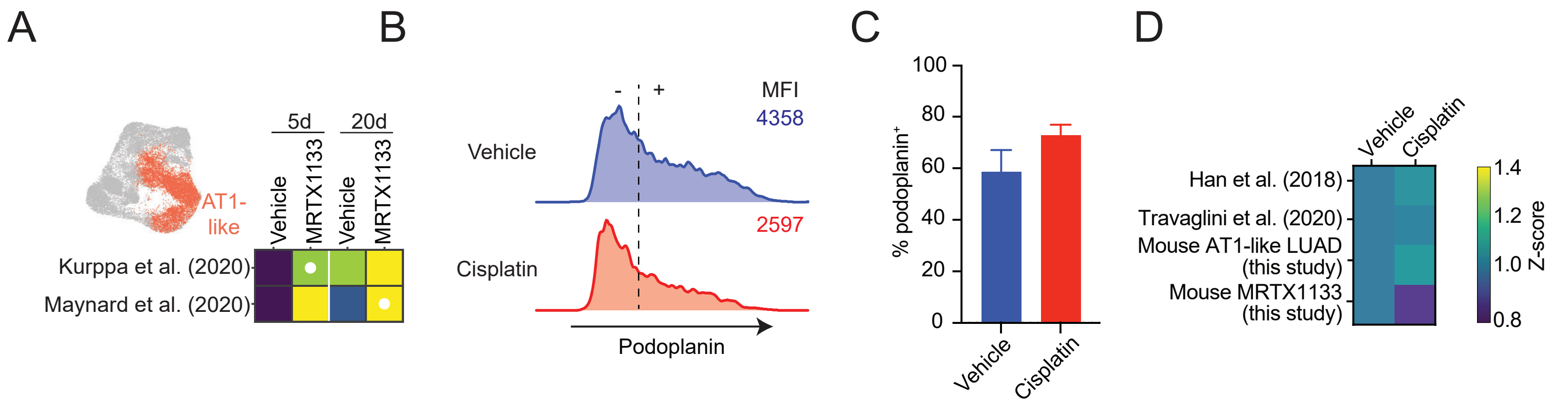


Li, Zhuang et al.
Supplementary Figure 3

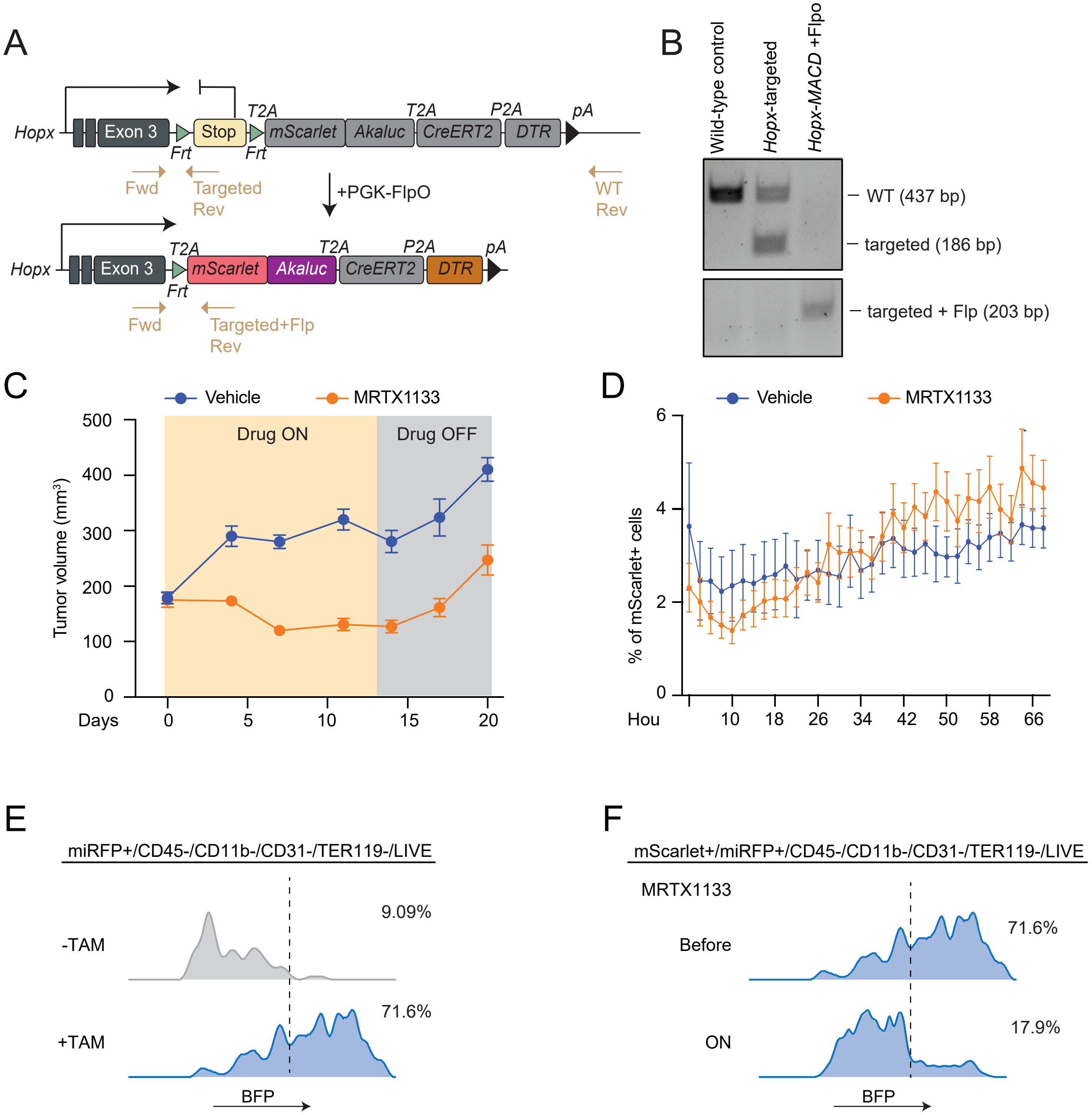


Li, Zhuang et al.
Supplementary Figure 4

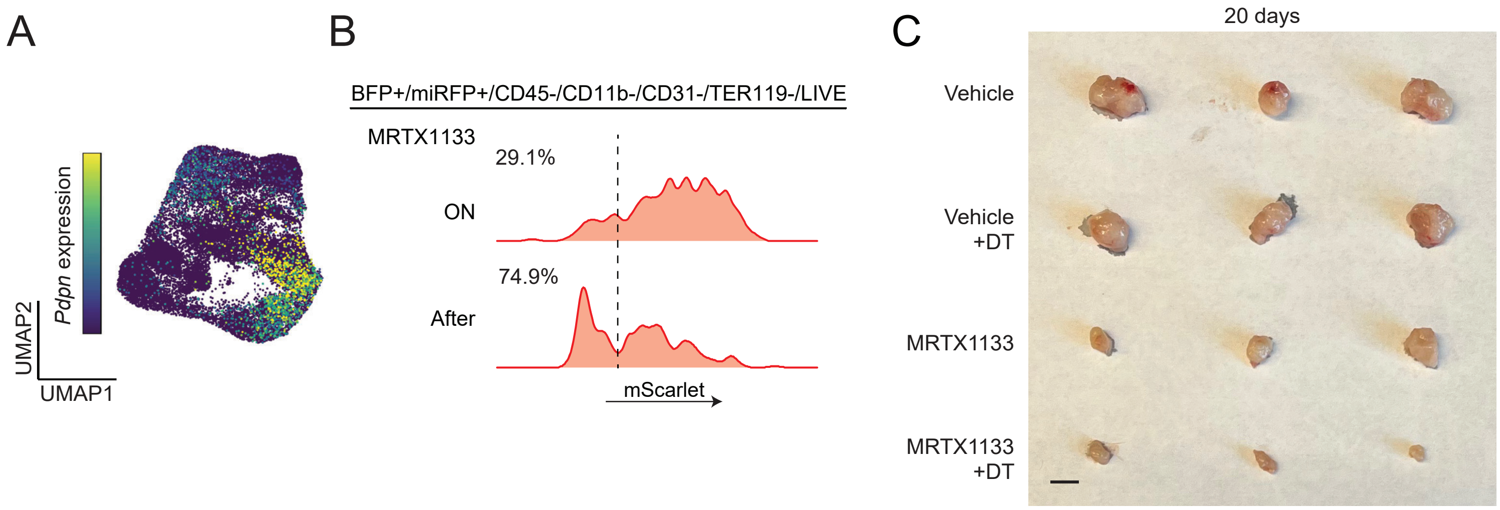


Li, Zhuang et al.
Supplementary Figure 5

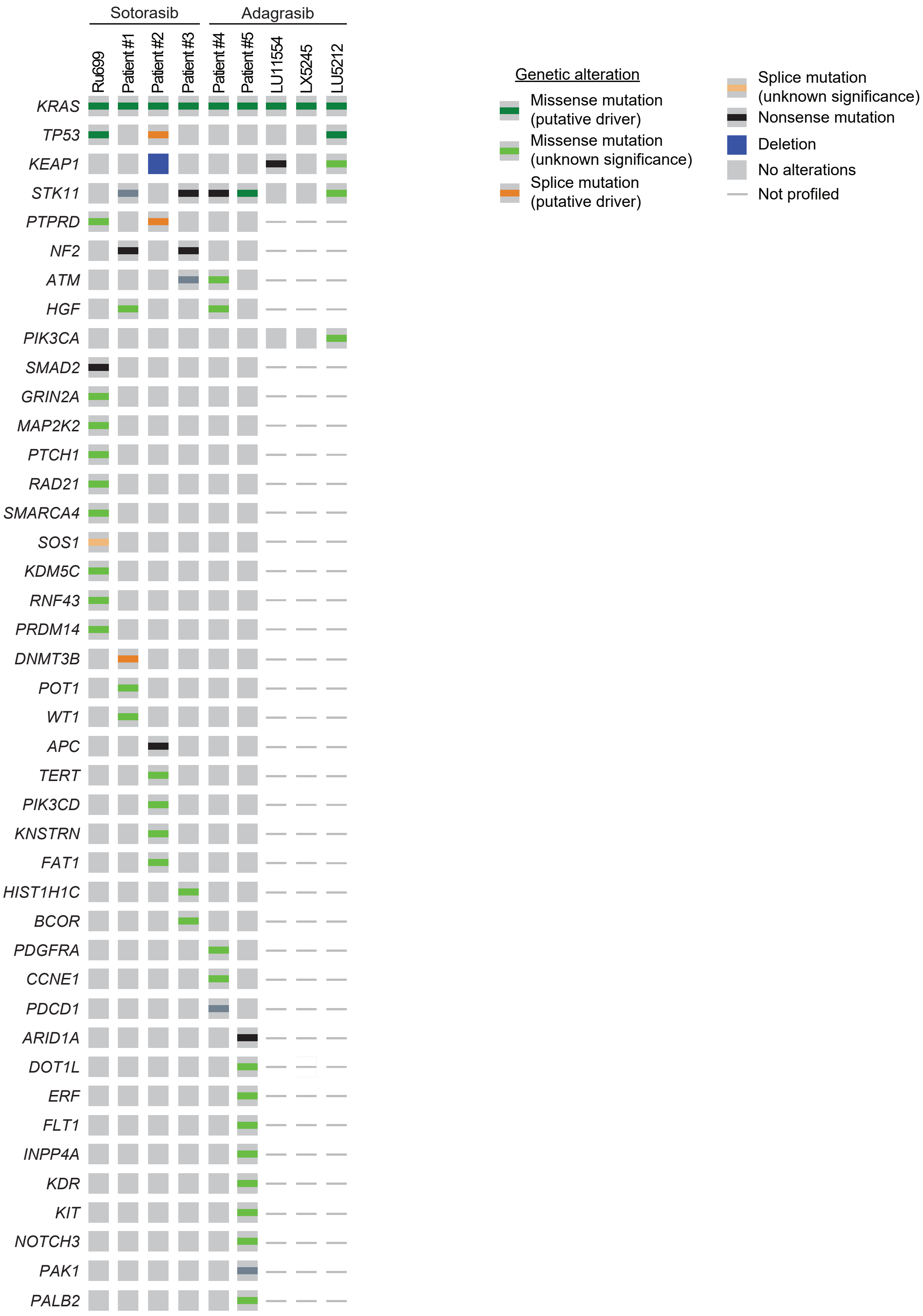


Li, Zhuang et al.
Supplementary Figure 6

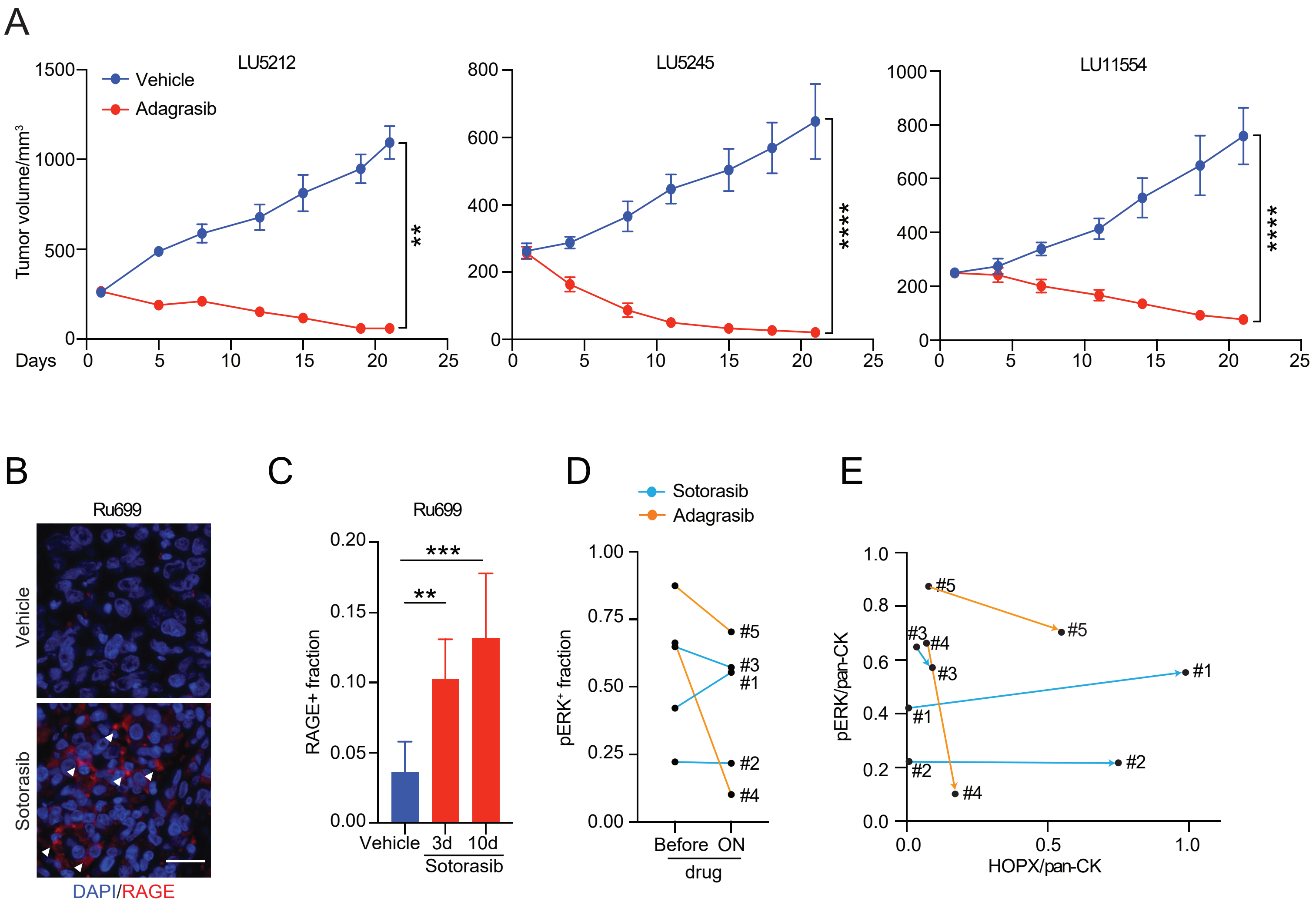


Li, Zhuang et al.
Supplementary Figure 7
